# Targeting Allostery in the Dynein Motor Domain with Small Molecule Inhibitors

**DOI:** 10.1101/2020.09.22.308700

**Authors:** Cristina C. Santarossa, Keith J. Mickolajczyk, Jonathan B. Steinman, Linas Urnavicius, Nan Chen, Yasuhiro Hirata, Yoshiyuki Fukase, Nicolas Coudray, Damian C. Ekiert, Gira Bhabha, Tarun M. Kapoor

## Abstract

Cytoplasmic dyneins are AAA (ATPase associated with diverse cellular activities) motor proteins responsible for microtubule minus-end-directed intracellular transport. Dynein’s unusually large size, four distinct nucleotide-binding sites, and the existence of closely-related isoforms with different functions, pose challenges for the design of potent and selective chemical inhibitors. Here we use structural approaches to develop a model for the inhibition of a well-characterized *S. cerevisiae* dynein construct by pyrazolo-pyrimidinone-based compounds. These data, along with single molecule experiments and mutagenesis studies, indicate that the compounds likely inhibit dynein by engaging the regulatory ATPase sites in the AAA3 and AAA4 domains, and not by interacting with dynein’s main catalytic site in the AAA1 domain. A double Walker B mutant in AAA3 and AAA4 is an inactive enzyme, suggesting that inhibiting these regulatory sites can have a similar effect to inhibiting AAA1. Our findings reveal how chemical inhibitors can be designed to disrupt allosteric communication across dynein’s AAA domains.

---

Dyneins are microtubule-based motor proteins that belong to the AAA (ATPase Associated with diverse cellular Activities) superfamily, which is defined by a widely conserved AAA domain (Erzberger and Berger, 2006). These motor proteins are conserved across many eukaryotes and are involved in a wide range of biological functions. In *Saccharomyces cerevisiae*, only one dynein isoform exists, and is required for the proper positioning of the nucleus during division (Moore et al., 2009). In metazoan cells there are two isoforms of cytoplasmic dynein that are responsible for microtubule minus-end directed transport. Dynein 1 is present in the cytoplasm and transports diverse cellular cargoes, including organelles, vesicles, chromosomes and mRNAs. In contrast, dynein 2 is localized to the cilium, where it drives retrograde intraflagellar transport (Roberts, 2018). As these transport processes occur on the timescale of seconds to minutes (Mijalkovic et al., 2017; Steinman et al., 2017), fast-acting small molecule probes can be powerful tools to examine dynein function with the required temporal resolution. Chemical inhibitors selective for dynein isoforms or targeting specific dynein-mediated protein interactions can also be useful therapeutics. For example, the life-cycle of viruses (e.g. HIV) in infected cells depends on dynein 1-mediated transport towards the nucleus of the host cell (Dharan and Campbell, 2018; Johnson et al., 1996; Pomeroy et al., 2002) and certain signal transduction pathways (e.g. Hedgehog signaling) needed for cancer cell growth depend on dynein 2 (Dharan and Campbell, 2018; Johnson et al., 1996; Pomeroy et al., 2002). However, we currently lack a framework for the design of potent and selective inhibitors of dynein.

At least three different cell-permeable inhibitors of dynein have been reported (Steinman and Kapoor, 2019). Ciliobrevins, the first cell permeable probes of dynein, were discovered using cell-based screens (Firestone et al., 2012). Dynapyrazoles, designed based on the ciliobrevin scaffold, have improved potency and inhibit the microtubule-stimulated, but not the basal, ATPase activity of human dynein (Steinman et al., 2017). Dynarrestin, which was also discovered using cell-based screens, can pull-down dynein complexes from cell extracts but does not inhibit ATPase activity *in vitro* (Höing et al., 2018). Structural models are needed to gain insights into the mechanism of dynein inhibition and to develop new chemical inhibitors, but these data have not been available.

Dynein’s motor domain, which powers microtubule-based motility, consists of a single polypeptide with six distinct AAA domains that form a ring, an N-terminal linker, and a long coiled-coil stalk that includes the microtubule binding domain (MTBD) (Figure 1A) (Carter et al., 2011; Kon et al., 2012). Another structural element, named the buttress (or strut), emanates from the AAA5 domain and interacts with the stalk, which extends from the AAA4 domain, to modulate the microtubule-binding affinity of the MTBD (Carter et al., 2011; Kon et al., 2012). Similar to other proteins in this superfamily, dynein’s six AAA domains are comprised of AAA-L (large) and AAA-S (small) subdomains that together with a AAA-L subdomain of an adjacent AAA domain, form a tripartite interface where ATP binding and hydrolysis can occur (Erzberger and Berger, 2006). The first four AAA domains (AAA1-AAA4) contain the Walker A motifs necessary for ATP binding, while the AAA5 and AAA6 domains lack this motif and likely have nucleotide-independent structural roles (Schmidt and Carter, 2016). The AAA1 domain contains the primary ATPase site that drives dynein’s motility (Gibbons and Gibbons, 1987; Kon et al., 2004). The AAA2 domain lacks the catalytic glutamate residue in the Walker B motif needed for ATP hydrolysis, resulting in constitutive ATP binding (Kon et al., 2012; Schmidt et al., 2012). The AAA3 active site plays an allosteric role, regulating ATP turnover in the AAA1 domain (Bhabha et al., 2014; DeWitt et al., 2015). The AAA4 domain contains the residues required for ATP binding and hydrolysis, but its function is not well understood (Kon et al., 2012; Schmidt et al., 2012). In principle, chemical inhibitors could bind at any of these four different AAA domains or another site in dynein to block its mechanochemical cycle.

**Figure 1.**
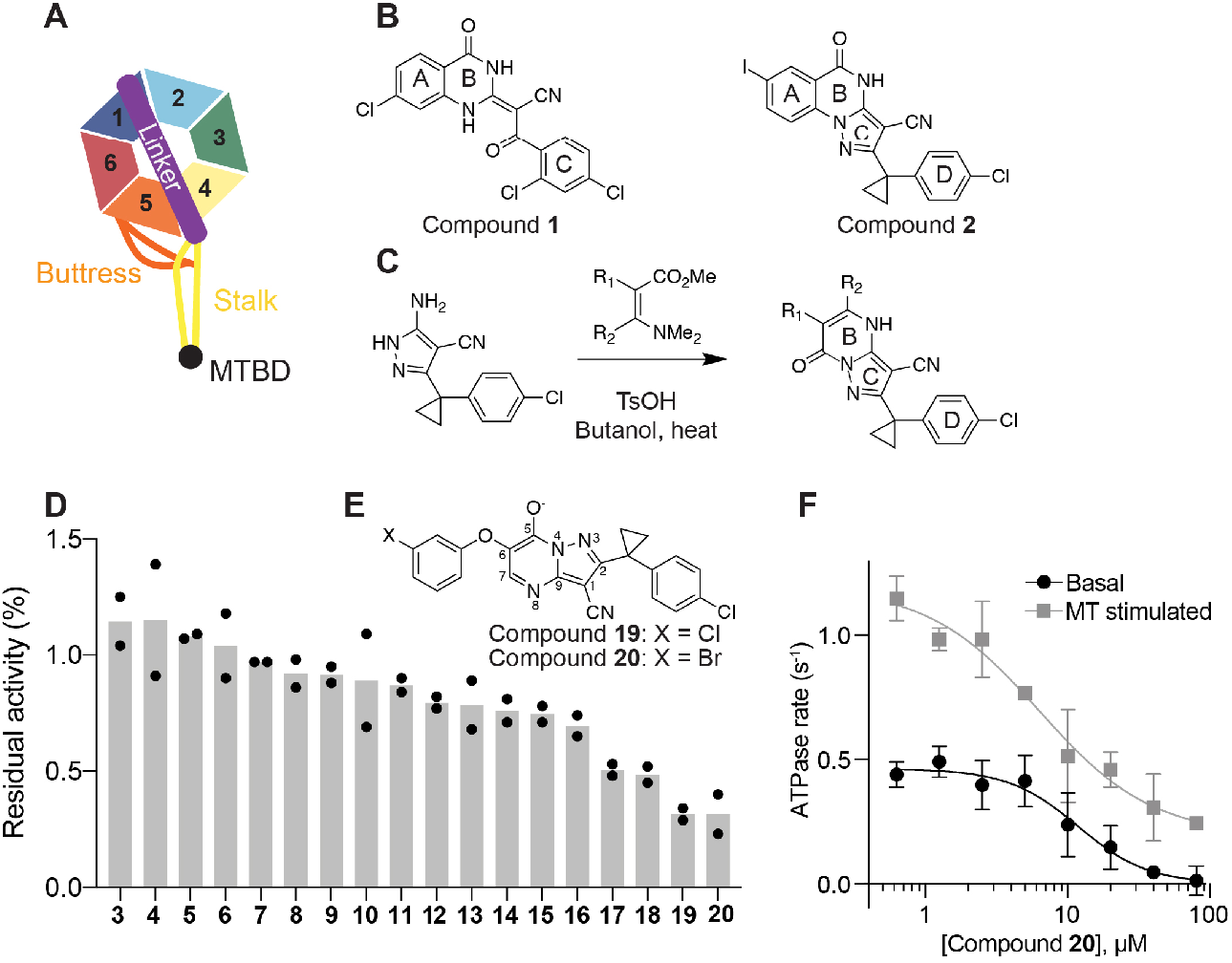
Inhibition of human dynein 1’s ATPase activity by dynapyrazole derivatives. (A) Schematic of dynein motor domain (AAA1: blue, AAA2: light blue, AAA3: green, AAA4: yellow, AAA5: orange, AAA6: red, linker: purple, buttress: orange, stalk: yellow, MTBD: black). (B) Chemical structure of the ciliobrevin D (compound 1) and dynapyrazole-A (compound 2). A, B, C and D rings are highlighted. (C) Synthesis of bicyclic dynapyrazole derivatives from a common precursor (TsOH: *p*-toluenelsulfonic acid). (D) Effect of dynapyrazole derivatives on Hs-dynein 1’s (N-terminal hexahistixine tag, aa. 1320–4646, uniprot: Q14204) basal ATPase activity. Compounds were tested at 20 μM. Data are mean ± range (n=2, for chemical structures see Figure S1A). (E) Chemical structures of compounds 19 and 20. Atom numbering is included. (F) ATPase activity of Hs-dynein 1 in the presence of compound 20 with or without the addition of microtubules (2.5 μM). Data are mean ± SD of n=3 and were fit to a sigmoidal dose-response curve.

Here we synthesized and tested derivatives of dynapyrazoles to identify inhibitors of the basal ATPase activity of dynein. We used X-ray crystallography, cryo-EM and computational docking to develop models for how a dynapyrazole derivative binds to *S. cerevisiae* dynein, which has been extensively used for structural studies of this motor protein (Bhabha et al., 2014; Carter et al., 2011; Niekamp et al., 2019; Schmidt et al., 2012). Single molecule fluorescence analyses were used to test the structural model for dynein inhibition. Our findings reveal how chemical inhibitors can disrupt the allosteric communication within dynein’s AAA ring and how two regulatory AAA sites contribute to enzymatic activity.

## Results

### Identifying dynapyrazole derivatives that inhibit dynein’s basal ATPase activity

To identify inhibitors of dynein’s basal ATPase activity we generated compounds that retained the pyrimidine-4-one based core that is common to the ciliobrevins and dynapyrazoles (hereafter, compound **1** and **2**) and therefore would likely maintain dynein inhibition (ring B, Figure 1B) (Firestone et al., 2012; See et al., 2016). A facile synthesis of these compounds was afforded by condensation of the 3-cyclopropyl-cholorophenyl substituted aminopyrazole with either 2- or 3-substituted (3-dimethylamino)acrylates in the presence of an acid to yield 5- or 6-substituted pyrazolo-pyrimidinones (Figure 1C).

We generated compounds **3-20** and tested their activity against a previously reported recombinant motor domain construct of human cytoplasmic dynein 1 (hereafter, Hs-dynein 1, aa. 1320-4646, Figures 1D and S1A-B) (Steinman et al., 2017). Hs-dynein 1’s basal ATP hydrolysis rate was 0.50 ± 0.13 s^-1^ (1 mM MgATP, mean ± standard deviation (SD), n>3, Figure 1F), consistent with previous studies (Steinman et al., 2017). Gratifyingly, we found that compounds (20 μM) with aryl-ether substitutions at the 6-position of the pyrazolopyrimidine core inhibited Hs-dynein 1’s ATPase activity in the absence of microtubules (Figure S1A). In particular, a *meta*-chlorophenoxy and *meta*-bromophenoxy ether substitution at the 6-position (compound **19** and **20**, respectively) led to the strongest inhibition (n=2, 1 mM MgATP, Figures 1D, 1E and S1A). Compounds with a *para*- or *ortho-chloro* groups on the pendent phenyl ring (compounds **17** and **13**, respectively) were less potent, as was an analog (**16**) with a *meta*-methoxy substitution (Figures 1D and S1A). Dose-dependent analyses indicated that compounds **19** and **20** inhibit Hs-dynein 1’s basal ATPase activity with an IC_50_ of 17 ± 2.3 μM and 12 ± 9.3 μM, respectively (1 mM MgATP, mean ± SD, n=3, Figures 1F, S1C).

We next measured the effect of compounds **19** and **20** on the microtubule-stimulated ATPase activity of Hs-dynein 1. In the presence of microtubules (2.5 μM), Hs-dynein 1’s ATPase rate was 1.02 ± 0.12 s^-1^ (mean ± SD, n>3), ~2-fold higher than the basal rate, as expected for this construct (Figures 1F, S1C) (Steinman et al., 2017). We note that tail-truncated dynein motor domain constructs have ATPase activities that are substantially lower than those reported for the multi-protein dynein-dynactin-BICD2 (DDB) complex, which can achieve rates of 100 s^-1^ (Cho et al., 2008; Elshenawy et al., 2019; Steinman et al., 2017). Dose-dependent analysis revealed that compounds **19** and **20** inhibited the microtubule-stimulated ATPase activity of Hs-dynein 1 with an estimated IC_50_ of ~30 and ~10 μM, respectively (1 mM MgATP, n=3, Figures 1F, S1C). Complete inhibition was not observed, possibly due to limited compound solubility of the inhibitors at high doses in this assay. We also examined if compounds **19** and **20** inhibit the ATPase activity of other AAA proteins that have been previously characterized: Hs-FIGL1, Mm-VCP, Xl-katanin, and Dm-spastin in the presence of compounds **19** and **20** (20 μM, 1 mM MgATP) (Cupido et al., 2019). These dynapyrazole analogs did not substantially inhibit (<25% reduction) of Mm-VCP, Xl-katanin, and Dm-spastin, but did inhibit Hs-FIGL1 (~85% inhibition) (n≥2, Figures S1D-E). Taken together, compounds **19** and **20** inhibit dynein’s basal ATPase activity, which compound **1** does not (Steinman et al., 2017). A compound which is capable of inhibiting the basal ATPase activity is more suitable for structural studies, as dynein constructs optimized for X-ray crystallography and cryo-EM can be utilized in the absence of microtubules. Therefore, compounds **19** and **20** facilitate structural work of dynein bound to a chemical inhibitor, and may serve as a scaffold for structure-guided design.

### Dynapyrazole derivatives inhibit *S. cerevisiae* dynein

One of the highest resolution (3.3 Å) structural models of the dynein motor domain was determined using *S. cerevisiae* dynein (Schmidt et al., 2012). Therefore, to maximize our chances of gaining structural insights into compound-motor domain interactions, we tested if compounds **19** and **20** also bound and inhibited *S. cerevisiae* dynein. For these experiments we first generated a previously described construct that contains the motor domain and has been characterized to be an active ATPase (VY137, aa. 1219-4093, hereafter, GFP-Sc-Dyn, Figures 2A, S1B) (Reck-Peterson et al., 2006). Dose-dependent analyses indicate that compounds **19** and **20** inhibit GFP-Sc-Dyn’s basal ATPase activity with an IC_50_ of 10 ± 1.2 and 14 ± 1 μM, respectively (1 mM MgATP, mean ± SD, n=3, Figure 2B).

**Figure 2.**
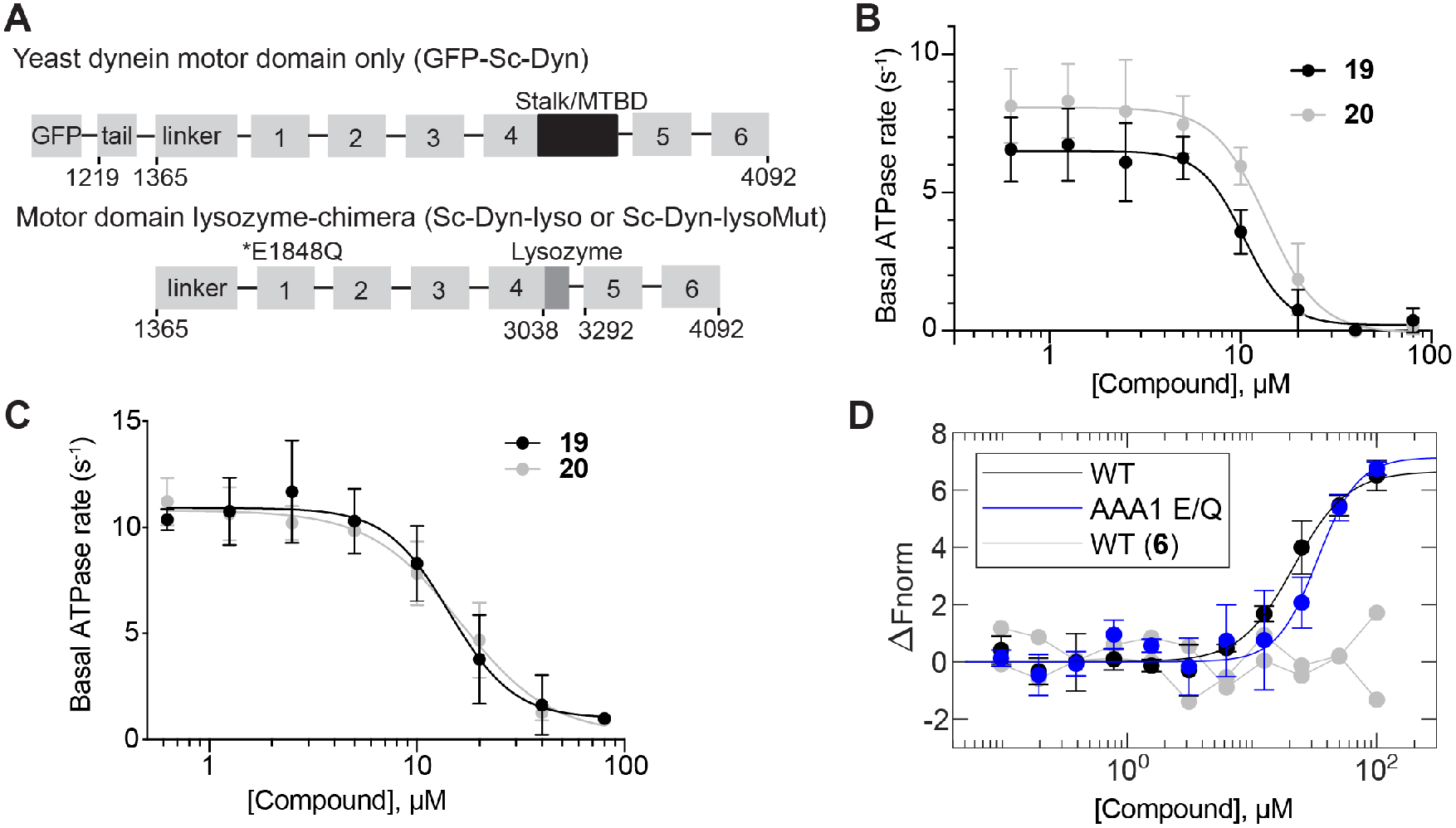
Dynapyrazole derivatives inhibit *Saccharomyces cerevisiae* dynein. (A) Schematic showing the domain organization for the *S. cerevisiae* dynein motor domain construct (GFP-Sc-Dyn) and either the WT lysozyme-chimera (Sc-Dyn-lyso) or lysozyme-chimera mutant constructs (Sc-Dyn-lysoMut). The position of the stalk and microtubule-binding domain (MTBD) in GFP-Sc-Dyn is shown (black box). A lysozyme replaces the MTBD and most of the stalk in the Sc-Dyn-lyso and Sc-Dyn-lysoMut constructs (dark gray box). The black asterisk indicates a Walker B mutation (E1849Q) in the AAA1 site of the Sc-Dyn-lysoMut construct. The first and last residues of each construct are indicated. (B) Basal ATPase activity of GFP-Sc-Dyn in the presence of compound **19** or **20**. Data are mean ± SD of n=3 and were fit to a sigmoidal dose-response curve. (C) Basal ATPase activity of Sc-Dyn-lyso in the presence of either compound **19** or **20**. Data are mean ± SD of n=3 and were fit to a sigmoidal dose-response curve. (D) Microscale thermophoresis analysis of compound **20**’s interactions with Sc-Dyn-lyso (WT) and Sc-Dyn-lysoMut (AAA1 E/Q). Compound **6** was used as a negative control. Data were fit using Equation (3) and converted to ΔF_norm_ using Equation (4) (see methods).

*S. cerevisiae* constructs that contain a Walker B mutation in the AAA1 site (E1849Q) and in which the microtubule binding domain (MTBD) and most of the coiled-coil stalk are replaced with lysozyme (VY972, hereafter, Sc-Dyn-lysoMut) have been shown to be well-suited for structural studies using X-ray crystallography and cryo-EM (Figures 2A, S1B) (Bhabha et al., 2014). As this construct is inactive, we used a construct with a wildtype, hydrolysis-competent AAA1 domain (E1849) that is otherwise identical to Sc-Dyn-lsyoMut (VY1027, hereafter, Sc-Dyn-lyso) (Figures 2A, S1B) and determined that compounds **19** and **20** inhibit Sc-Dyn-lyso’s ATPase activity with an IC_50_ of 14 ± 4.9 μM and 15 ± 3.1 μM, respectively (1 mM MgATP, mean ± SD, n=3, Figure 2C).

For structural studies, we focused on compound **20**, as bromine’s anomalous diffraction may aid in locating the compound in the X-ray diffraction data (Arkhipova et al., 2017). To examine binding of compound **20** to Sc-Dyn-lyso and Sc-Dyn-lysoMut, we used microscale thermophoresis (MST), a technique that monitors temperature-induced changes in the mobility of fluorescent molecules as a readout to measure intermolecular interactions (Jerabek-Willemsen et al., 2014). Compound **20** binds to Sc-Dyn-lyso and Sc-Dyn-lysoMut with an estimated EC_50_ of 22 ± 5.3 μM and 34 ± 11 μM, respectively (mean ± 95% confidence interval (CI), n=3, Figure 2D). As a negative control, we also tested compound **6,** which does not inhibit the ATPase activity of Hs-dynein 1 at the 20 μM dose (Figures 1D, S1A). The assay did not reveal binding of compound **6** to Sc-Dyn-lyso, consistent with our ATPase data (n=2, Figure 2D). These results indicate that a Walker B mutation at the AAA1 site in dynein does not affect compound **20** binding. Taken together, these data suggest that the Sc-Dyn-lysoMut construct can be employed for our structural studies in the presence of compound **20**.

### X-ray model of *S. cerevisiae* dynein (Sc-Dyn-lysoMut) in the presence of a dynapyrazole derivative

We obtained diffracting crystals of Sc-Dyn-lysoMut in the presence of compound **20** and in the absence of nucleotide. Our screening conditions were based on those previously reported for this construct in the presence of AMPPNP (Bhabha et al., 2014). We solved the structure by molecular replacement to ~4.5 Å resolution (Figures 3A, 3B and Table S1). At this resolution most of the α helices are well resolved but many of the individual β strands are not (Figure S2A). Density for each of the six AAA domains, as well as the linker, truncated stalk and lysozyme is observed (Figure 3B).

**Figure 3.**
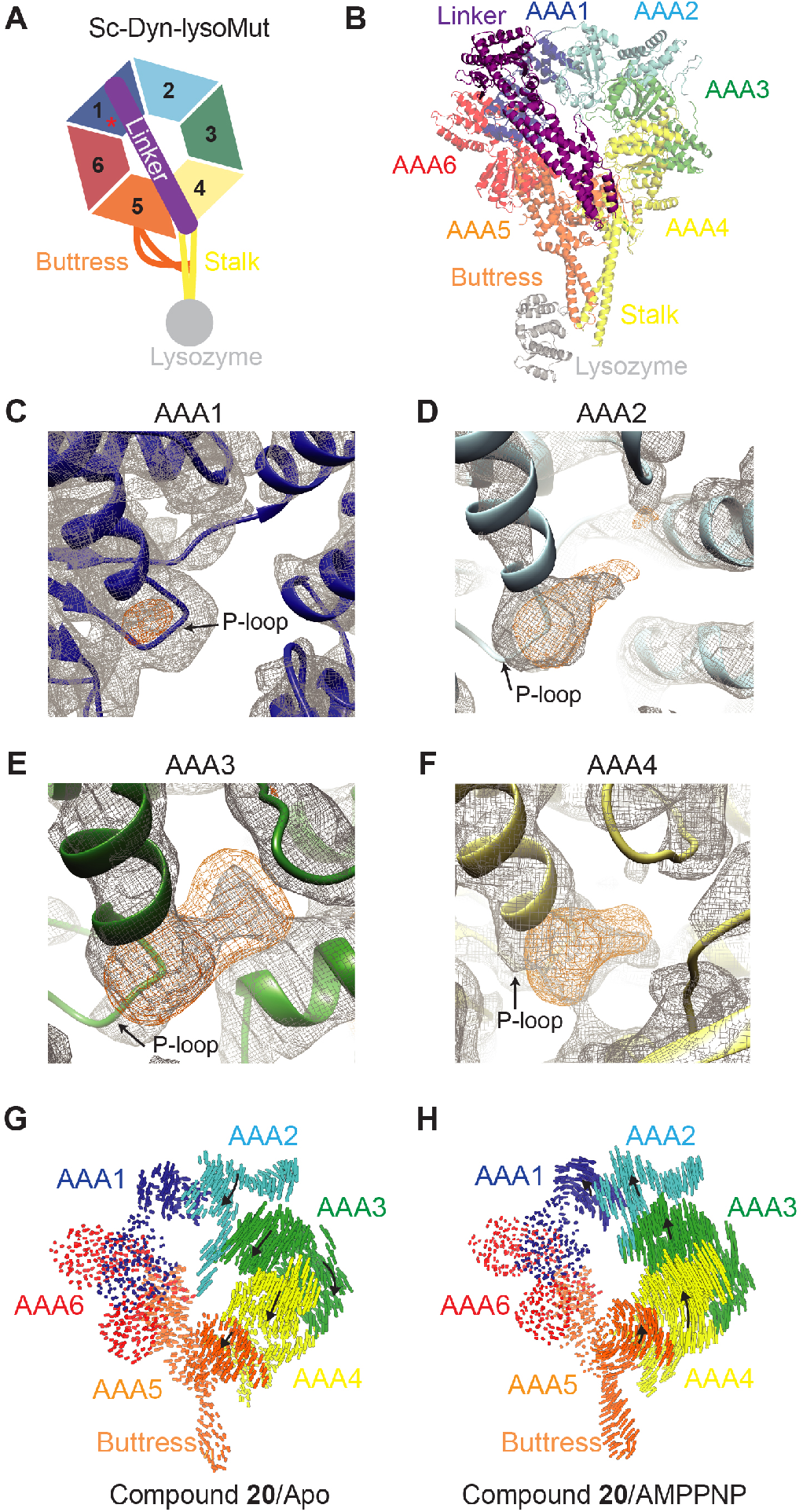
X-ray model of Sc-Dyn-lysoMut in the presence of compound 20. (A) Schematic of the Sc-Dyn-lysoMut construct. Individual AAA domains are shown (AAA1: blue, AAA2: light blue, AAA3: green, AAA4: yellow, AAA5: orange, AAA6: red). Linker (purple), buttress (orange), stalk (yellow) and lysozyme (gray) are also depicted. Red asterisk indicates that the construct contains a Walker B mutation (E1848Q) in the AAA1 domain. (B) Refined model of Sc-Dyn-lysoMut. Domains are color-coded based on the schematic shown in Figure 3A. (C-F) *2Fo – Fc* (gray mesh, 1.0 σ contour) and *Fo* – *Fc* (orange mesh, 3.0 σ contour) densities are shown in the AAA1 (C), AAA2 (D), AAA3 (E), and AAA4 (F) domains. (G, H) Visualization of interalpha carbon distances between the X-ray model and either the apo (PDB: 4AKG) (G) or AMPPNP (PDB: 4W8F) model (H). Models were aligned on the AAA1-L subdomain. Linker was removed for clarity. Black arrows indicate direction of domain movement in the X-ray model relative to either of the two nucleotide states, while the size of the arrow indicates the magnitude of movement.

We first analyzed electron density in the nucleotide-binding pockets that are not accounted for by the protein backbone. In the AAA1 domain, a small positive electron density is observed near the P-loop in the Fo-Fc map (Figure 3C). Based on its overall size, this density may correspond to an anion (Figure 3C). In the AAA2, AAA3 and AAA4 nucleotide-binding sites, electron densities are present in both the 2Fo-Fc and Fo-Fc maps (Figures 3D–3F). These densities are larger than what would be expected for an anion or water molecule and may correspond to either nucleotide or compound **20**, but are not well-resolved enough to interpret. In addition, an anomalous signal was not detected in the diffraction data, possibly due to the flexibility of the bromo-phenyl moiety, low compound affinity, or the current resolution. Overall, hints of additional density were observed in the AAA2, AAA3 and AAA4 nucleotide-binding sites, but establishing their identity was not possible due to the resolution of this model.

We next examined the conformation of the individual AAA domains. The AAA1 domain is in an open conformation, similar to what is observed in the nucleotide-free model of *S. cerevisiae* dynein (pdb 4AKG, hereafter, apo-model) (Figures S2B-S2D), consistent with no nucleotide or chemical inhibitor bound. In the apo-model, the AAA3 domain adopts a semi-closed conformation relative to the AAA1 domain, which can close further upon nucleotide binding (Bhabha et al., 2014; Kon et al., 2012; Schmidt et al., 2012; Zhang et al., 2017). The conformation of the AAA3 domain in our X-ray model is similar to that of the apo-model (Figures S2E-G), suggesting that nucleotide is not bound to this site in our crystal.

We also examined the overall conformation of the AAA ring in the presence of compound **20**. We observed that the linker is extended and docked onto the AAA5-L subdomain (Figure 3B), indicating that the motor domain is in a post-powerstroke state (Burgess et al., 2003). Therefore, we aligned the Sc-Dyn-lysoMut X-ray model (hereafter, X-ray model) to other structural models of *S. cerevisiae* dynein with a similarly extended linker conformation such as those that represent the nucleotide-free (apo-model) and AMPPNP-bound states (pdb 4W8F, hereafter AMPPNP-model) (Bhabha et al., 2014; Schmidt et al., 2012). Alignment of the X-ray model with the apo-model on the AAA1-L subdomain revealed that the stalk, buttress and the AAA5-AAA6 domains adopt similar conformations (Figure 3G). However, the AAA2/AAA3/AAA4 domains, as a unit, rotate (~8°) towards the linker, when compared to the apo-model. A similar alignment showed that the AAA2/AAA3/AAA4 unit has rotated much less in the X-ray model compared to the AMPPNP-model (Figure 3H). The AAA ring is more planar in the X-ray model compared to the apo-model, but less planar than that in the AMPPNP-model (Figures S2H, S2I, and S2J). Taken together, the X-ray model reveals a conformational change in one-half of the AAA ring, despite the lack of nucleotide or chemical inhibitor density in the AAA1 site. Interestingly, this conformation has only been observed in structural models of dynein where AMPPNP is bound to the AAA1 site (Bhabha et al., 2014; Niekamp et al., 2019). However, a higher resolution model of Sc-Dyn-lysoMut in the presence of compound **20** is needed to determine if the inhibitor binds to the AAA3 or AAA4 sites.

### Cryo-EM model of *S. cerevisiae* dynein (Sc-Dyn-lysoMut) in the presence of compound 20

Our X-ray model clearly revealed the conformation of the AAA ring in the presence of compound **20**, but the resolution did not allow us to identify the binding site(s) of the chemical inhibitor. Therefore, we next turned to single-particle cryo-EM studies, using the same mutant construct (Sc-Dyn-lysoMut) we used for X-ray crystallography (Bhabha et al., 2014). We hypothesized that a large cryo-EM dataset may lead to a higher resolution model for the motor protein, as we would have the opportunity to classify compositional and conformational heterogeneity, which may be limiting the resolution of our X-ray structure.

We prepared samples of Sc-Dyn-lysoMut in the presence of compound **20** (80 μM) and without any nucleotide present, and collected a large dataset on a Titan Krios microscope equipped with K2 camera (4892 movies, Figures S3, S4A, and S4B). Due to the challenges presented by sample heterogeneity and particle damage during the freezing process, we used extensive data processing and classification strategies, which included data from two other datasets in the initial processing steps (described in detail in methods and Figure S3). After 3D classification and refinement we obtained a reconstruction with an average resolution of ~3.9 Å with 136,180 particles (Map 1, Figures S3B and S4C). We found that the overall conformation of dynein’s AAA ring in this reconstruction was similar to that observed in our X-ray model (Figure 4A), as indicated by the root-mean square displacement (RMSD) of 1.512 Å (2354 c-alpha positions).

**Figure 4.**
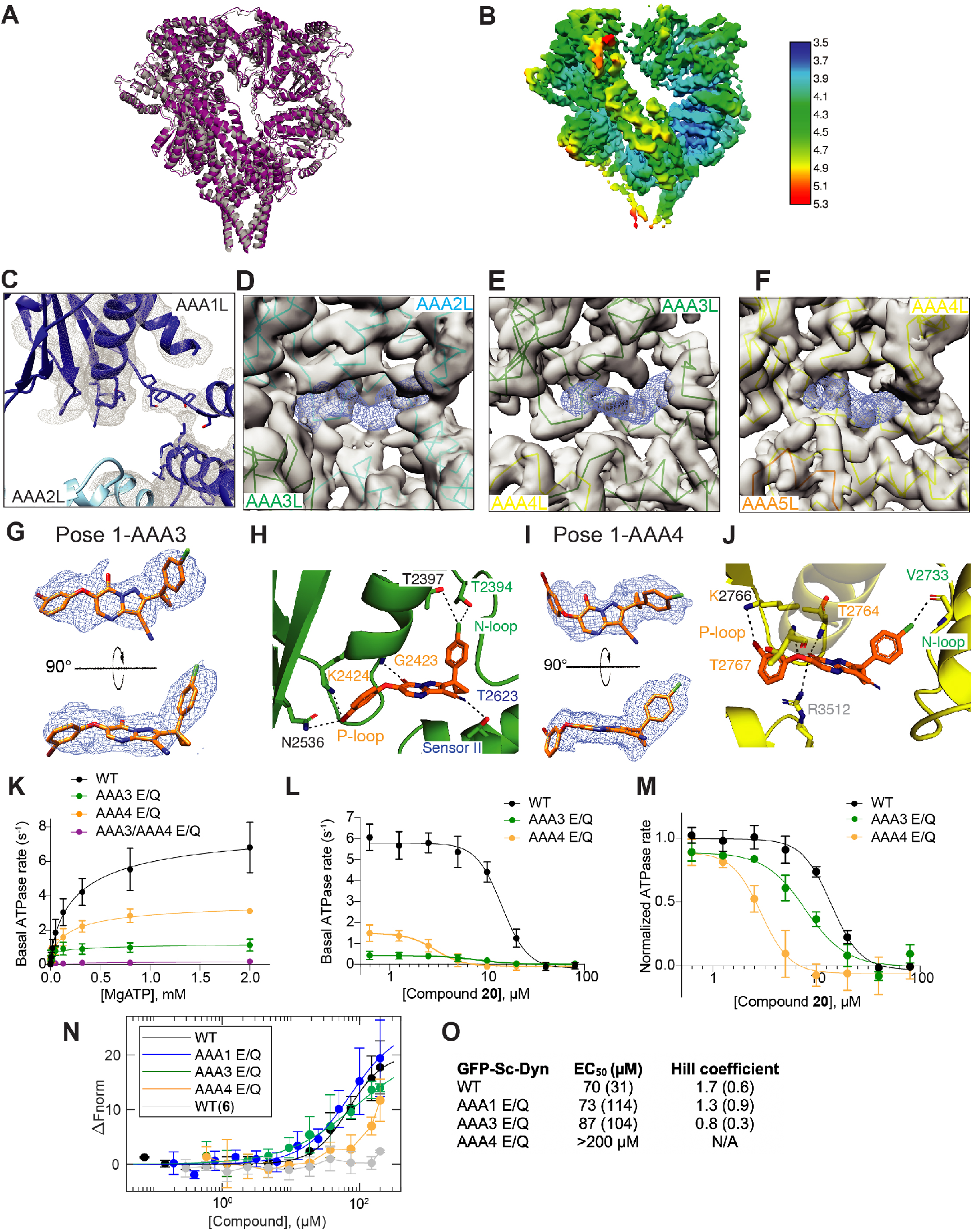
Cryo-EM model of Sc-Dyn-lysoMut in the presence of compound 20. (A) X-ray model (gray) aligned to the cryo-EM model of Sc-Dyn-lysoMut (purple). Lysozyme is omitted from the X-ray model for calculation of RMSD between the two models. (B) EM map (Map 1) of Sc-Dyn-lysoMut colored by local resolution, as estimated by RELION 3.0. (C-E) EM density corresponding to the nucleotide-binding sites of either the AAA1 (Map 2) (C), AAA2 (Map 4) (D), AAA3 (Map 4) (E), or AAA4 (Map 4) (F) domains. Densities (Map 4) corresponding to the Sc-Dyn-lysoMut model (AAA2L: cyan, AAA3L: green, AAA4L: yellow, AAA5L: orange) are shown in gray, whereas densities (Map 4) for the ligand are shown as blue meshes. For each AAA domain, the location of the large and small subdomains are indicated. (G) Two views of pose 1-AAA3 overlayed with the EM density (Map 4, blue mesh). Compound **20** is shown as a stick model (carbon: orange, oxygen: red, nitrogen: blue, chlorine: green, bromine: dark red). (H) View of compound **20** in the nucleotide-binding site of the AAA3 domain. The predicted inhibitor-protein interactions (black dashed line) are shown (N-loop: green, P-loop: yellow, sensor II: blue) (I) Two views of pose 1-AAA4 overlayed with the EM density (Map 4, blue mesh). Compound **20** is shown as a stick model (carbon: orange, oxygen: red, nitrogen: blue, chlorine: green, bromine: dark red). (J) View of compound **20** in the nucleotide-binding site of the AAA4 domain. The predicted inhibitor-protein interactions (black dashed line) are shown (N-loop: green, P-loop: yellow, sensor II: blue, arginine finger: gray). (K) ATP-concentration dependence of the steady-state activity of GFP-Sc-Dyn and its mutants, analyzed using an NADH-coupled assay. Rates were fit to the Michaelis-Menten equation for cooperative enzymes (mean ± SD, n=3, Equation 2). (L) Concentration-dependent inhibition of the steady-state ATPase activity of GFP-Sc-Dyn and its mutants by compound **20**. Data were fit using a sigmoidal dose response equation (Equation 1). AAA3 E/Q and AAA4 E/Q mutants had an IC_50_ of 8.2 ± 1.4 μM and 2.9 ± 0.3 μM, respectively (mean ± SD, n=3). (M) Data from panel L replotted as rate relative to DMSO control. (N) Microscale thermophoresis analyses of GFP-Sc-Dyn and its mutants in the presence of either compound **6** or **20**. Data are shown as mean ± SD and fitted using the Hill equation (n>3, Equation 3). Fits were weighted by the inverse standard error of the mean at each data point. (O) EC_50_ values and hill coefficients are shown for each construct (fitted value, with 95% CI in parentheses).

Based on the differences in local resolution (Figure 4B), it was clear that heterogeneity within the AAA ring was still limiting the resolution of the EM map. Therefore we used masking, signal subtraction, and 3D classification to process densities corresponding to the AAA2-AAA4, AAA5-AAA6 and AAA6-AAA2 domains separately, resulting in three independent reconstructions (see methods, Figure S3B). This process revealed heterogeneity in the AAA1 domain, such that density for the AAA1-S subdomain is either present (Map 2, ~4.5 Å) or weak/missing (Map 3, ~7.9 Å). When density for the AAA1 domain is clearly defined, we can conclude that AAA1 is in an open state, as expected when it is nucleotide-free. After further masking and signal subtraction, we obtained higher resolution maps of the AAA2/AAA3/AAA4 (Map 4, ~3.5 Å) and AAA5/AAA6 domains (Map 5, ~3.7 Å) (Figures S3B, S4D and Table S2). Consistent with these resolutions, densities for many side chains are observed (Figure S4E).

The cryo-EM map revealed no additional density in the AAA1 nucleotide-binding pocket, (Figure 4C), which is consistent with the observation that the AAA1 domain adopts an open conformation. However, the AAA2, AAA3 and AAA4 nucleotide-binding pockets all clearly show the presence of additional density that is not accounted for by the protein (Figure 4D–4F). AAA2 lacks catalytic activity and is therefore expected to be constitutively bound to ATP (Schmidt et al., 2012). Indeed, the map reveals a ligand density in the AAA2 nucleotide-binding site that is consistent with the size and shape of ATP (Figure S5A).

We next examined the cryo-EM map that corresponds to the AAA3 and AAA4 binding sites, which revealed additional densities that are consistent with the size and shape of compound **20** (Figures 4G and 4I). Three lines of evidence suggest that this density corresponds to compound **20**. First, the AAA3 domain is in a semi-closed conformation, similar to the X-ray- and apo-models, and it is therefore unlikely that nucleotide is bound at the AAA3 site (Figure S5D). Second, the protein was purified in the absence of nucleotide, which is unlikely to yield a protein containing ATP or ADP in the AAA1, AAA3 and AAA4 sites, as previously shown (Schmidt et al., 2012). Third, if nucleotide were present, we would expect it to be bound at AAA1, especially since the E1849Q mutation renders this domain hydrolysis deficient and would retain nucleotide in the binding site. Therefore, we posit that the densities in the AAA3 and AAA4 sites correspond to compound **20**, which was present in 50-fold molar excess in our sample relative to the protein concentration.

### Computational docking of compound 20 into the proposed inhibitor binding sites

The local resolution of our cryo-EM reconstruction at the inhibitor binding sites is not high enough to unambiguously identify the inhibitor binding pose. Therefore, we performed computational docking of compound **20** into the AAA3 and AAA4 sites and ATP into the AAA2 site using the GlideEM script in the Schrodinger software (Schrodinger LLC) (see methods) (Robertson et al., 2019). This script utilizes the GLIDE docking scoring function and the real space cross-correlation score to the EM map to generate binding poses of a ligand that are energetically favored and that can be accommodated by the density (Robertson et al., 2019). The enol tautomer of compound **20** was used for computational docking, which was calculated to be the major tautomer at pH 7 (see methods). This analysis generated five candidate poses of compound **20** for the AAA3 site, and four poses for the AAA4 site (Figures 4G, 4I, S5B, S5C) (Robertson et al., 2019). For the AAA2 site, the GlideEM script revealed a pose of ATP that is similar to that previously reported (Schmidt et al., 2012). For the AAA3 and AAA4 sites, we selected the orientation of the ligand that best fits into the EM density, as indicated by the docking model-EM map correlation coefficient (Afonine et al., 2018). The poses that best fit into the EM densities in the AAA3 and AAA4 sites (hereafter, pose 1-AAA3 and pose 1-AAA4) had a value of 0.73 and 0.74, respectively, indicating a good fit (Figures 4G, 4I, S5B and S5C) (Afonine et al., 2018). Visual inspection of the map also clearly identified this as an appropriate binding pose.

For pose 1-AAA3, the 3-cyclopropyl-chlorophenyl moiety of compound **20** is buried in the adenine-binding pocket and is within van der waals distance to hydrophobic residues in the N-terminal loop motif (N-loop) (Figures 4H and S5E). The backbone of the phosphate-binding loop (P-loop) residue G2423, which interacts with the β-phosphate of AMPPNP in the AMPPNP-model (chain B), forms a hydrogen-bonding interaction with the carbonyl oxygen of the enolatepyrimidinone moiety (Figures 4H and S5E) (Bhabha et al., 2014). The N-loop residue, T2394, and a residue that neighbors the N-loop motif, T2397, form a halogen-bonding interaction with the chloro-phenyl moiety. The threonine in the sensor II motif (T2623) hydrogen bonds with the cyano moiety, while the catalytic lysine (K2424) and residue N2536 form halogen bonds with the bromo-phenyl group. These proposed interactions may explain how compound **20** can displace nucleotide to bind to the AAA3 ATP-binding site.

In pose 1-AAA4, compound **20** binds to the nucleotide-binding pocket in the AAA4 domain in a similar orientation shown for the AAA3 site (Figures 4J and S5F). The 3-cyclopropyl phenyl moiety fits into the adenine-binding pocket of the AAA4 site and is stabilized by hydrophobic interactions with N-loop residues V2730, P2731, and V2733. The chloro-phenyl moiety forms a halogen-bonding interaction with the backbone of N-loop residue V2733. Additionally, the arginine finger (R3512) and the backbone of P-loop residue, T2764, hydrogen bond with the carbonyl oxygen of the enolatepyrimidinone moiety. The backbone of P-loop residue T2767 forms a hydrogen-bonding interaction with the phenoxy moiety of compound **20**. Taken together, these analyses suggest that compound **20** can adopt a similar orientation in both the AAA3 and AAA4 sites and is capable of forming a network of interactions with the P- and N-loops motifs in dynein.

### Mutagenesis studies are consistent with compound 20 binding to the AAA3 and AAA4 domains

Our structure suggests that compound **20** binds to the regulatory ATPase sites in the AAA3 and AAA4 domains. This is an intriguing finding given that inhibitors of dynein’s ATPase activity would be predicted to interact with the main catalytic site in AAA1. Our structural model suggests that simultaneously inhibiting AAA3 and AAA4 allosterically inhibits the enzyme, and we would then hypothesize that mutations inhibiting ATP hydrolysis in AAA3 and AAA4 would result in similarly low ATPase activity as a mutation inhibiting hydrolysis in AAA1. To test this hypothesis, we used GFP-Sc-Dyn mutants containing the following Walker B mutations: AAA3 E/Q (E2488Q), AAA4 E/Q (E2819Q), and AAA3/AAA4 E/Q (E2488Q/E2819Q) (Figure S1B). The basal catalytic turnover number (k_cat_) of GFP-Sc-Dyn and its mutants was measured using a steady-state ATPase assay (Figures 4K, S5G). GFP-Sc-Dyn had a k_cat_ of ~8 s^-1^, which is within 2-fold of the published value (~4 s^-1^) for an *S. cervisiae* motor domain construct of dynein (n=5, Figures 4K, S5G, S5I) (Cho et al., 2008; DeWitt et al., 2015). The k_cat_ values of the AAA3 E/Q and AAA4 E/Q mutants were reduced by ~6- and ~2-fold, respectively, relative to that of the GFP-Sc-Dyn, consistent with published results (Figures 4K, S5G, S5I) (Cho et al., 2008; DeWitt et al., 2015). Notably, Walker B mutations in both the AAA3 and AAA4 sites yielded an inactive enzyme, with ATPase rates near the baseline, similar to inhibition reported for the AAA1 E/Q mutation (Nicholas et al., 2015). These results suggest blocking ATP hydrolysis in the AAA3 and AAA4 sites is a plausible mechanism to inhibit dynein’s basal ATPase activity.

We next examined if the Walker B mutations in the AAA3 and AAA4 sites affect the potency of compound **20** in the steady-state ATPase assay. Compound **20** inhibited GFP-Sc-Dyn with an IC_50_ of 14 ± 0.8 μM (1 mM MgATP, mean ± SD, n=3, Figures 4L-M). The AAA3 E/Q and AAA4 E/Q mutants were more sensitive to compound **20** inhibition, with IC_50_ values ~2- and ~5-fold lower than that of GFP-Sc-Dyn, respectively (1 mM MgATP, n=3, Figures 4L-M). We note that compound **20**’s shift in potency was not due to a decrease in the mutants’ affinity for ATP, as indicated by the K1/2 (Figure S5H, S5I). These data show that mutations in the AAA3 and AAA4 sites affect the sensitivity of dynein to compound **20** inhibition.

Since the ATPase assay is a combined readout of ATP hydrolysis in the AAA1, AAA3, and AAA4 sites, the effect of the Walker B mutations on compound **20**’s activity is not a direct indicator of binding. We therefore measured the binding of compound **20** to GFP-Sc-Dyn using MST. Compound **20** bound to GFP-Sc-Dyn with an EC_50_ of 70 ± 31 μM and a hill slope of 1.7 ± 0.6, indicating positive cooperativity (mean ± 95% CI, n=9, Figures 4N-O). Complete saturation of the binding curve could not be obtained, likely due to limited compound solubility in this assay. As a negative control, we also tested compound **6,** which did not appear to bind to GFP-Sc-Dyn in this assay. We note that the GFP-Sc-Dyn EC_50_ is higher than that observed for Sc-Dyn-lyso, possibly due to technical differences in the MST assay, where Sc-Dyn-lyso and its mutant are labeled with an amine-reactive dye while the GFP tag of GFP-Sc-Dyn is used as the fluorescent readout (see methods). Nonetheless, compound **20** inhibited the ATPase activities of GFP-Sc-Dyn and Sc-Dyn-lyso with similar potencies, suggesting that the two constructs behave similarly with respect to inhibition by compound **20** (Figures 2B and 2C).

We next tested the AAA1 E/Q (E1849Q) mutant, which we generated as a negative control since our structures suggest that the compound does not bind to AAA1 (Figure S1B). Consistent with our structural data, the binding of compound **20** to the AAA1 E/Q mutant is similar to WT, with an EC_50_ of 73 ± 114 μM and a hill slope of 1.3 ± 0.9 (mean ± 95% Cl, n=4, Figures 4N-O). While the AAA3 E/Q mutant bound to compound **20** mutant with a similar EC_50_ as the WT protein (EC_50_: 87 ± 104, mean ± 95% Cl, n=4), we observed a significant reduction in the hill slope relative to that of GFP-Sc-Dyn (h: 0.8 ± 0.3, mean ± 95% Cl, n=4), indicating loss of cooperativity (Figures 4N-O). Notably, compound **20** binding to the AAA4 E/Q mutant was substantially decreased compared to the WT curve (EC_50_: >200 μM, n=4, Figures 4N-O). Taken together, ATP inhibition and binding assays data are consistent with our structural model, where compound **20** interacts with the AAA3 and AAA4 domains, and suggest that inhibiting these two domains simultaneously can inhibit the enzyme in similar manner as would be expected when inhibiting the main catalytic site, AAA1.

### Compound 20 inhibits the dynein landing rate in single-molecule assays

To test the effect of compound **20** on dynein motility, we performed single-molecule motility assays using total internal reflection fluorescence (TIRF) microscopy (Figure 5A). For these studies we used an N-terminal GFP-tagged GST-dimerized motor (VY208; hereafter, Sc-DynGST), as the motility parameters for this dynein construct have already been characterized (Bhabha et al., 2014; Cho et al., 2008; DeWitt et al., 2015; Reck-Peterson et al., 2006). We measured the landing rate, velocity, and processivity (run length) of Sc-DynGST motors on rhodamine-labeled, taxol-stabilized microtubules over a range of compound **20** concentrations.

**Figure 5.**
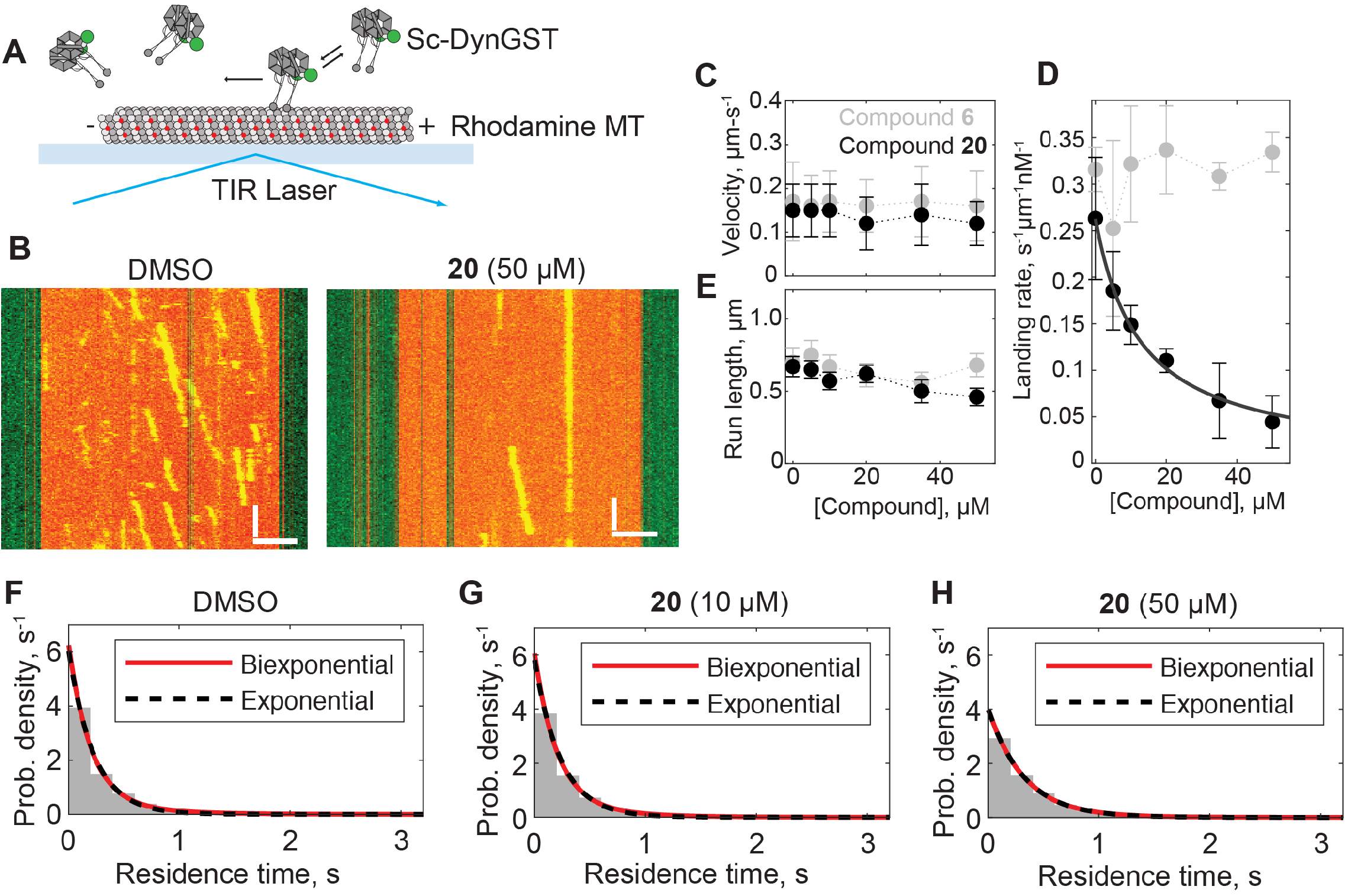
Using single molecule assays to analyze the inhibition of *S. cerevisiae* dynein by compound 20. (A) Schematic of microtubule motility assay. Sc-DynGST, rhodamine-labeled microtubule (MT), coverslide (light blue) and total internal reflection fluorescence (TIRF) laser are indicated. (B) Example kymographs of Sc-DynGST in the presence of 1% DMSO or compound **20** (50 μM). Kymograph horizontal scale bar, 2.5 μM; kymograph vertical scale bar, 5 s. (C) Analysis of Sc-DynGST landing rates on microtubules in the presence of either compound **6** or **20**. For compound **6**, data are shown as mean ± SD for three independent experiments containing a total of 657-794 motile events from 78 microtubules in each condition. Dotted gray line is to guide the eye. For compound **20**, data are shown as mean ± SD for n ≥ 3 independent experiments containing a total of 219-1097 motile events from 58-86 microtubules, and fitted to a sigmoidal dose-response equation. (D-E) Velocities (D) and run lengths (E) of Sc-DynGST motors at various concentrations of either compound **6** or **20**. Velocities determined as sample mean ± SD and run lengths determined as fit ± bootstrapped error using an offset exponential fit to the cumulative density function for n=210-259 measurements for compound **6** and n=193-390 measurements for compound **20** (data pooled from 3-4 independent experiments). Entire distributions are shown in Figure S6 and S7. Dotted lines are shown to guide the eye. (H) Probability distribution of residence times of individual Sc-DynGST molecules within 50-nm bins along the direction of motion, following a published method (DeWitt et al., 2015). Here significant pausing should yield a second, slower, exponential phase. Fitting to single exponentials (black dashed lines) and biexponentials (red lines) yielded nearly identical results. Data are also shown in Figure S6.

We found that compound **20** induces a dose-dependent decrease in the landing rate, or the number of new dynein motility events per unit time per unit microtubule length per nM dynein (Figures 5B and 5C). The EC_50_ for the dynein landing rate was 12.7 ± 2.5 μM (fit ± 95% confidence intervals), which is within error of the IC_50_ for Sc-Dyn-lyso’s basal ATPase activity in the presence of compound **20** (Figures 2C and 5C). Hence, the presence of compound **20** reduces dynein’s ability to bind microtubules.

The presence of compound **20** had little to no effect on dynein’s velocity or processivity (Figures 5D, 5E and S6), once a dynein molecule had landed on a microtubule. As ATP hydrolysis at AAA1 is required for each step that dynein takes, we would expect either a reduction in velocity or processivity if compound **20** bound to this site (Bhabha et al., 2014; Gibbons and Gibbons, 1987; Kon et al., 2004). We also observed no significant pausing during processive runs (Figures 5F–5H and S6), as has been seen when AAA3 is locked in an ATP-like state (DeWitt et al., 2015). As a compound control, we repeated the single-molecule assay with compound **6**. The landing rate, velocity, and run length of Sc-DynGST were not sensitive to this compound (Figures 5C-E and S7). Together, these data are consistent with a model in which compound **20** binds to the regulatory ATPase sites, rather than the main catalytic site in the AAA1 domain.

## Discussion

While inhibitors of dynein have been described before, their binding sites have been difficult to identify. This is largely due to the inherent flexibility of the dynein motor domain, which makes it a challenging target for structural biology and limits achievable resolution. Here we describe derivatives of dynapyrazoles that inhibit the ATPase activity of human dynein 1 and *S. cerevisiae* dynein, and report X-ray and cryo-EM models of the dynein motor domain in the presence of compound **20**. The resolution of our cryo-EM model is comparable to that of prior X-ray models of the *S. cerevisiae* dynein motor domain (3.3-3.8 Å) (Bhabha et al., 2014; Schmidt et al., 2012), and is substantially higher than those reported for other cryo-EM models of the dynein motor domain (~7-20 Å) (Bhabha et al., 2014; Niekamp et al., 2019; Toropova et al., 2014). This model reveals the inhibitor binding sites, as density for compound **20** is observed in the AAA3 and AAA4 domains. Functional single molecule motility assays show that in the presence of compound **20**, velocity and processivity, which are dependent on ATP binding and hydrolysis at the AAA1 site, are not affected, suggesting that compound **20** does not bind AAA1. Together, our findings suggest that compound **20** inhibits dynein’s ATPase activity by disrupting allosteric communication across the AAA ring.

Our binding data suggests that compound **20** interacts with the AAA3 and AAA4 domains. However, mutating the Walker B motif in the two domains resulted in different effects on compound **20** binding. For the AAA3 E/Q mutant, the EC_50_ was similar to that of the WT protein, but the hill slope was significantly reduced. For the AAA4 E/Q mutant, the EC_50_ was substantially increased. The environment of the AAA3 and AAA4 Walker B residues are likely different, as previously suggested (Schmidt et al., 2012), which may explain the differences in the binding data. While future work and higher resolution is needed to probe exactly how compound **20** binds to the AAA3 and AAA4 domains, three lines of biochemical evidence suggest that the inhibitor targets these sites. First, the MST data shows that the AAA3 E2488Q mutation results in loss of cooperativity, suggesting that ligand binding reduces the affinity of dynein to compound **20**. The binding affinity of AAA4 E2819Q mutation to compound was substantially reduced. Second, loss of ATP binding or hydrolysis in either AAA3 or AAA4 is not sufficient to inactivate dynein’s ATPase activity (Cho et al., 2008; Kon et al., 2004), indicating that both sites need to be inhibited, since the main catalytic site in AAA1 does not appear to interact with compound **20**. Third, the ATPase inhibition data by the dynapyrazole derivative suggests that mutations in the AAA3 and AAA4 domain are capable of affecting compound activity. Thus, our findings reveal that mutations in the AAA3 and AAA4 sites are able to affect dynein’s affinity and sensitivity to compound **20**.

The dynapyrazole derivative binds to the AAA3 and AAA4 sites, resulting in a conformational change in the AAA ring relative to both the apo- and AMPPNP-model. Current models suggest that ATP binding to the AAA1 site initiates a rigid body movement of the AAA2/AAA3/AAA4 unit towards the linker domain (Bhabha et al., 2014; Schmidt et al., 2015). These conformational changes are propagated to the other half of the ring (AAA5/AAA6), which in turn pulls the buttress motif relative to the stalk domain to alter microtubule binding affinity (Figure 6) (Bhabha et al., 2014; Niekamp et al., 2019; Schmidt et al., 2015). In our model, compound **20** binding to the AAA3 and AAA4 sites results in a rotation of the AAA2/AAA3/AAA4 block towards the linker domain to produce a more planar AAA ring, a conformation that is intermediate between the apo and AMPPNP-bound states. Similar conformational changes were observed in a model of a dynein stalk mutant solved in the presence of AMPPNP (Niekamp et al., 2019). This stalk mutation has been proposed to decouple the two halves of the AAA ring, resulting in weak microtubule binding affinity. We propose that compound binding to the AAA3 and AAA4 sites similarly interfere with the propagation of conformational changes between the two halves of the AAA ring, also resulting in weak microtubule attachment (Figure 6).

**Figure 6.**
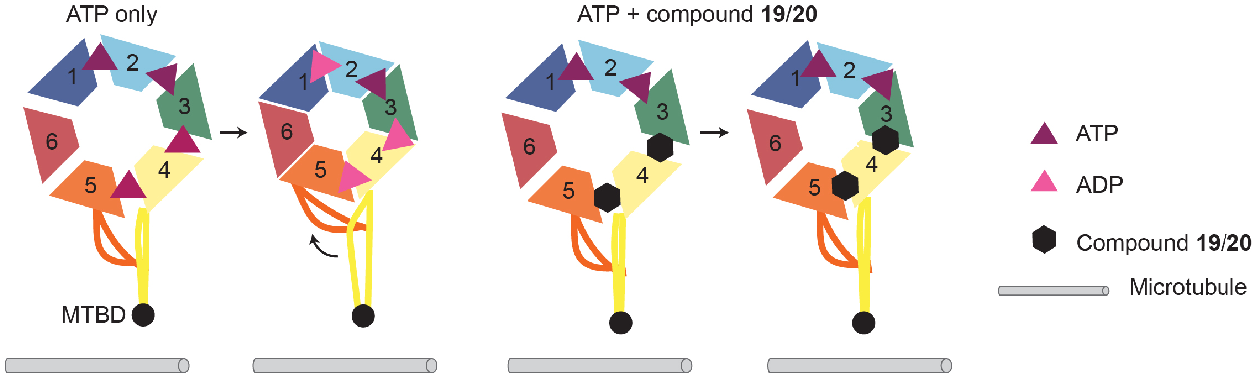
Model for the inhibition of *S. cerevisiae* dynein by the dynapyrazole derivatives. Schematic of the *S. cerevisiae* dynein motor domain. Individual domains are shown and color-coded based on the schematic shown in Figure 3A. The microtubule binding domain (MTBD) at the tip of the stalk is shown (black circle). Upon ATP binding and hydrolysis in the AAA1, AAA3, and AAA4 domains, the two halves of the AAA ring rotate towards each other, resulting in closure of the AAA ring, and the buttress slides relative to the stalk, changing the stalk registry. In the presence of ATP (purple triangle) and compound 20 (black hexagon), the nucleotide binds to the AAA1 site, but cannot be hydrolyzed. One-half (AAA2-AAA4) of AAA ring rotates, but is not sufficient to induce conformational changes in the other half of the ring. The buttress does not slide relative to the stalk, resulting in weak microtubule binding affinity.

Computational docking suggests key interactions with ATP-binding residues in the AAA3 and AAA4 sites that likely contribute to dynein inhibition by these compounds. Compound **20** adopts an ATP-like conformation, where the cyclopropyl-phenyl moiety buries itself within the adenine-binding site, and the halo-phenoxy enolatepyrimidinone moieties interact with the P-loop motif. This model may explain why compounds with phenyl-ether substitutions at the 6-position, specifically with small substituents at the *para* or *ortho* positions (i.e. compounds **17-20**), more potently inhibit human dynein’s ATPase activity (≤50% residual activity at 20 μM, Figures 1D, S1A) since they can more likely adopt an ATP-like conformation compared to the inactive analogs. Notably, the proposed inhibitor-target interactions are different from those observed for other active-site binding chemical inhibitors of AAA proteins, such as spastazoline and CB-5083, which contact residues in the N-loop and sensor II motifs (Pisa et al., 2019; Tang et al., 2019). Further studies will examine if modifying the scaffold to allow for hydrogen-bonding interactions with the N-loop motif can lead to an improvement in potency.

Dynein’s motor domain is a multi-site enzyme with a complex mechanism of allostery that can be leveraged for the development of new inhibitors. We show that the ATP-binding pockets of the AAA3 and AAA4 domains are druggable allosteric sites that can be targeted to inhibit the main catalytic activity in AAA1. This mode of inhibition is in contrast to allosteric inhibitors that have been described for other AAA proteins, such as VCP/p97 and VPS4, which bind ~20 Å away from the ATPase site (Banerjee et al., 2016; Pöhler et al., 2018). Thus, our data reveal a new strategy to inhibit *S. cerevisaie* dynein’s ATPase activity, and future work will investigate whether this mechanism applies to different isoforms of human dynein, such as dynein 2.

## Acknowledgements

T.M.K. is grateful to the NIH/NIGMS for funding (GM98579 and R35 GM130234-01). G.B. is grateful to the NIH/NIGMS R00GM112982 and the Damon Runyon Cancer Research Foundation DFS-20-16 for funding. C.C.S. was supported by the Tri-Institutional PhD Program in Chemical Biology and the Rockefeller University Graduate Program. K.J.M. was supported by a National Cancer Institute K00 Fellowship (K00CA223018). We thank Tommaso Cupido and Rudolf Pisa for providing us the following recombinant AAA proteins: Dm-spastin, Xl-katanin, Mm-VCP and Hs-FIGL1. We are especially grateful to Ronald Vale and the Vale lab for the gift of the VY137, VY972, VY1027, VY208 and VY696 *S. cerevisiae* strains. We thank James Fraser (University of California, San Francisco), Gydo van Zundert (Schrodinger LLC), and Kenneth Borrelli (Schrodinger LLC) for help with computational docking. We thank Carolina Adura at The Rockefeller University High-Throughput and Spectroscopy Resource Center and Deena Oren at The Rockefeller University Structural Biology Resource Center for assistance. The use of the Formulator robot in the Rockefeller University Structural Biology Resource Center was made possible by 1S10RR027037-01 from the National Center for Research Resources of the NIH. We also thank Mark Ebrahim and Johanna Sotiris at the Evelyn Gruss Lipper CryoEM Resource Center at The Rockefeller University for assistance in data collection. This research used the 17-ID-2 beamline of the National Synchrotron Light Source II, a US Department of Energy (DOE) Office of Science User Facility operated by the Brookhaven National Laboratory under contract no. DE-SC0012704.

## Authors contributions

C.C.S., G.B, and T.M.K. conceived the project and designed experiments. C.C.S. synthesized and tested compounds, designed mutant constructs, purified proteins, performed cryo-EM and X-ray experiments, processed data, built molecular models, and performed microscale thermophoresis experiments. K.J.M. performed the single-molecule assays and analyses. J.B.S. initially identified compound **19** as an inhibitor of human dynein 1. L.U. and N.C. assisted with cryo-EM data collection and processing. N.C., Y.H. and Y.F. synthesized the dynapyrazole derivatives. D.C.E. assisted with X-ray data collection and processing. C.C.S, G.B., and T.M.K. prepared the manuscript.

## Methods

### Protein purification

The human cytoplasmic dynein 1 construct (Hs-Dynein 1, uniprot: Q14204) containing an N-terminal hexahistixine (6x-His) was expressed using the baculovirus/insect cell expression system and purified as previously described (Steinman et al., 2017). *Xenopus laevis* katanin (Xl-katanin), *Mus musculus* VCP (Mm-VCP), *Homo sapiens* FIGL1 (Hs-FIGL1), and *Drosophila melanogaster* spastin (Dm-spastin) were expressed and purified as previously described (Cupido et al., 2019).

All *S. cerevisiae* dynein constructs were purified as previously described with some modifications (Reck-Peterson et al., 2006). Eight liters of *S. cerevisiae* cells were grown until stationary phase and harvested by centrifugation. Cells were washed with water, and resuspended in 5X lysis buffer (150 mM HEPES, pH 7.4, 250 mM K-Ac, 10 mM Mg(Ac)_2_, 2 mM EGTA). Cells were slowly pipetted into liquid nitrogen and then lysed using a coffee grinder. The resulting powder was resuspended in 2.5X lysis buffer (75 mM HEPES, pH 7.4, 125 mM K-Ac, 5 mM Mg(Ac)_2_, 0.5 mM EGTA, 3 mM DTT, 0.2 mM ATP, 1 mM PMSF) and clarified by ultracentrifugation (37,000 rpm for 2 hours). The supernatant was poured into a column containing IgG beads (Sepharose 6 Fast Flow, GE Healthcare) and allowed to bind to the beads by gravity-flow. The beads were washed with wash buffer (30 mM HEPES, pH 7.4, 50 mM K-Ac, 2 mM Mg-Ac, 1 mM EGTA, 10% glycerol, 200 mM KCl, 2 mM DTT, 100 μM ATP, 0.4 mM PMSF, 0.25% Triton-X). Beads were subsequently washed with TEV buffer (50 mM Tris-HCl, pH 7.4, 150 mM K-Ac, 2 mM Mg(Ac)_2_, 1 mM EGTA, 10% glycerol, 0.5 M TCEP, 0.1 M PMSF) and resuspended in TEV buffer (3 mL) containing TEV protease (25 μg) for overnight on-column TEV cleavage at 4°C. Protein was eluted and concentrated using an Amicon Ultra (100 kDa). The protein was subjected to size-exclusion chromatography (Superdex 200I, GE Healthcare) and eluted in size-exclusion buffer (20 mM Tris, pH 8.0, 50 mM K-Ac, 2 mM Mg(Ac)_2_, 1 mM EGTA, 10% glycerol, 1 mM TCEP).

### Co-crystallization of Sc-Dyn-lysoMut with compound 20

Sc-Dyn-lysoMut was concentrated to ~8 mg/mL and compound **20** was added (final compound concentration 100 μM, 2% DMSO) in size-exclusion buffer. A seed stock was prepared by using Seed Bead (Hampton Research, HR2-320) to crush crystals of Sc-Dyn-lysoMut that were generated using a previous condition (Bhabha et al., 2014). Seeds were added to the protein-compound complex (1:4 dilution) and mixed with an equal amount of reservoir solution (100 mM Bis-Tris pH 6.8-7.2, 200 mM sodium acetate, 8-13.5% PEG 3350, and 10 mM TCEP) using the sitting drop method. Crystals were allowed to form at 18°C. Prior to cooling in liquid nitrogen, the crystals were supplemented with ethylene glycol to give a final precipitant (PEG 3350, ethylene glycol) concentration of 35%.

### Data Collection and Refinement

Diffraction data were collected at the NSLS-II 17-ID-2 (FMX) beamline (wavelength: 0.920094 Å) at Brookhaven National Laboratory and indexed to P21212 and reduced using XDS (Table S1) (Kabsch, 2010). The X-ray data was phased by molecular replacement using Phaser (McCoy et al., 2007). A nucleotide-free model of the *S. cerevisiae* dynein motor domain (PDB: 4AI6) was used as the search model to yield a solution. The model was adjusted in Coot (Emsley et al., 2010) and refined using Phenix (McCoy et al., 2007). The lysozyme, which is not present in the search model, was built by rigid body docking of the lysozyme coordinates from PDB: 4W8F into the defined electron density. Ramachandran statistics are 94% favored, 6% allowed, 0%outliers.

### Cryo-EM sample preparation and data collection

For cryo-EM of *S. cerevisiae* (Sc-Dyn-lysoMut) in the presence of compound **20**, freshly purified protein was diluted to either 0.15 or 0.5 mg/mL and mixed with 2X compound **20** stock solution (final concentration: 80 μM compound **20**, 0.2% DMSO). Samples were applied to glow discharged Quantifoil 1.2/1.3 copper grids and plunge frozen in liquid ethane using Vitrobot IV (Thermo Fisher Scientific). Dose-fractionated (50 frames, 10s exposure, 44 e^-^ per Å^2^) super-resolution image stacks were collected on a 300-kV Titan Krios electron microscope (Thermo Fisher Scientific) equipped with a K2-summit detector (Gatan) using automated data collection (SerialEM). Datasets were collected over three separate sessions, where the first two were collected at 1.3 Å per pixel and the last collected at 1.0 Å per pixel.

### Data processing and analysis

Two datasets were originally acquired on two different grids that only differed by their protein concentration (0.15 vs 0.5 mg/ml). For these first two datasets, per pixel drift correction and dose-weighting was performed using MotionCor2. The motion-corrected, dose-weighted images were imported into CryoSPARC and contrast transfer function (CTF) parameters were estimated using CTFFind4. For dataset 1, approximately 1,000 particles were manually picked and subjected to 2D classification. Classes that represented different orientations of the protein were used as templates for automatic particle picking. Multiple rounds of 2D classification were performed to discard classes that displayed ice or carbon contamination, or poorly resolved averages. Particles from the classes with the best alignment parameters were selected and used to build an *ab initio* model as a reference for 3D refinement. A similar process was followed for dataset 2, but for the final refinement, the 124,891 particles selected were combined with the 25,016 particles from the dataset 1, reaching a resolution of ~4.4 Å. The results obtained from those datasets were subsequently used at three stages for the processing of dataset 3. The 2D classes were used to auto-pick, the best particles were then used as seeds to drive 2D classification, and the 3D map was used as a reference for 3D refinement.

Dataset 3 was collected in super-resolution at a higher pixel size (0.518 Å/pixel) and processed using RELION 3.0. First, correction of inter-frame movement for each pixel and dose-weighting was performed using RELION 3.0’s implementation of MotionCor2. Motion-corrected images were first binned to 1.3 Å per pixel and the CTF parameters were estimated as previously described (Rohou and Grigorieff, 2015). 2D class averages from the first two datasets were used as templates for automatic particle picking in RELION 3.0. To expedite 2D classification, the 4,723,853 particles picked were split into 24 sub-sets. Addition of the best 149,907 particles from datasets 1 and 2 to each subset facilitated the separation of good particles from junk during the 2D classification of dataset 3. Each subset was then subjected to seven rounds of 2D classification. Particles selected from dataset 3 were re-extracted at a pixel size of 0.518 A/pixel. While the 3D model obtained from dataset 2 was used as a reference, only particles from dataset 3 were used for refinement. After a round of 3D classification with eight classes, the two best classes were merged and refined to 3.9 Å.

Local resolution estimation revealed that the AAA3 and AAA4 domains contained the highest resolution information, while the map showed lack of density for the AAA1 domain. To increase local resolution, we performed a wide range of signal subtractions to refine individual or adjacent AAA domains. Among those, three locally refined maps of 2 or 3 adjacent AAA domains lead to improved resolution: AAA6-AAA1 (Map 2), AAA2-AAA4 (Map 4), and AAA5-AAA6 (Map 5).

The model was constructed using coordinates from the X-ray model presented here. Rigid body fit of each individual AAA small and large domain was performed using PHENIX, as part of the real-space refinement procedure (ADP refinement, local grid search, secondary structure restraints, Ramachandran restraints). Coot, Chimera, and Pymol were used to visualize and analyze the maps and models.

### Computational docking and refinement

Ligand docking was performed using the GlideEM script in the Schrödinger software (Robertson et al., 2019). The individual AAA2, AAA3, and AAA4 domains were extracted from the model and subjected to computational docking separately. The major tautomer at pH 7.0 of compound **20** was assigned performed using the LigPrep panel in Maestro. Default parameters were used for GlideEM docking, except for the following modifications: RECEP_CCUT=0.24,RECEP_VSCALE=0.9, EPIK_ PENALTIES=False, HBOND_DONOR_AROMH=True, HBOND_ACCEP_HALO=True. Real space refinement was performed in PHENIX using default settings except that the OPLS3e /VSGB2.1 force field was used with a weight factor of 1.

### Radioactive ATPase assay

ATPase assays using Hs-dynein 1 were performed as previously described, except that the protein concentration was reduced to 30 nM and the reaction time was extended to 45 min (Steinman et al., 2017). If microtubules were present, the buffer (25 mM PIPES pH7.0, 30 mM KCl, 1 mM EGTA, 5 mM MgCl2, 0.01% Triton™ X-100, 1 mM DTT) contained 20 μM taxol. For structure-activity relationship studies with the dynapyrazole derivatives, percent inhibition of the ATPase activity was calculated by normalizing the ATPase rate in the presence of compounds to DMSO control. To determine the IC_50_ of compound **20,** the measured activity was plotted against concentration of compound **20** and the data were fit using the sigmoidal dose response curve (Equation (1)) in Prism v. 8.0 (GraphPad Software Inc).

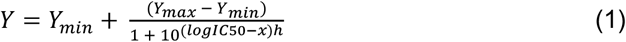

### NADH-coupled steady state ATPase assay for *k_cat_* analyses

To determine the *k*_cat_ and *k*_1/2_ of GFP-Sc-Dyn and its mutants, an enzymatic ATP regeneration system was used. For GFP-Sc-Dyn and its mutants, the assay buffer contains 30 mM HEPES pH 7.4, 50 mM K-Acetate, 2 mM magnesium acetate, 1 mM EGTA, 0.01% Triton X-100, and 1 mM DTT. A 20 μL reaction was prepared in a flat bottom 384 well black polystyrene plate (Greiner Bio One, Wemmel, Belgium, catalog # 781090) and contained protein (Sc-Dyn-lyso: 5 nM, GFP-Sc-Dyn: 50 nM) and 1X NADH mix. 1X NADH mix contains 200 μM NADH (Sigma, N7410), 1 mM phosphoenol pyruvic acid monopotassium salt (Sigma, P7127), 33.3 U/mL D-lactic dehydrogenase (Sigma, L3888), and 55 U/mL pyruvate kinase (lyophilized powder, Sigma, P9136). The reaction was initiated with the addition of 10X MgATP (2 μL), and the fluorescence signal was monitored using a Synergy NEO Microplate Reader (λex = 340 nm, 440 nm emission filter). The slope of the fluorescence values versus time was calculated to give the rate of fluorescence decrease. The ATPase rate was calculated from the rate of fluorescence decrease from a standard curve measuring the contribution of ADP to the NADH fluorescence signal. Enzyme parameters *k*_1/2_, *k*_cat_, and Hill coefficients (*h*) for the recombinant enzymes were determined by fitting the rates to the Hill equation using Prism v. 6.0 (GraphPad Software Inc) at different ATP concentrations (*x*):

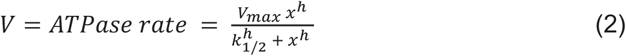

### NADH-coupled steady state ATPase assay for analysis of compound 19 and 20

A 20 μL reaction was prepared in a flat bottom 384 well black polystyrene plate (Greiner Bio One, Wemmel, Belgium, catalog # 781090) and contained Sc-Dyn-lyso, 1X NADH mix, and compound. 1X NADH mix contains 200 μM NADH (Sigma, N7410), 1 mM phosphoenol pyruvic acid monopotassium salt (Sigma, P7127), 33.3 U/mL D-lactic dehydrogenase (Sigma, L3888), and 55 U/mL pyruvate kinase (lyophilized powder, Sigma, P9136). For Sc-Dyn-lyso, a ten dose serial dilution was performed in DMSO starting at 4 mM compound **19** or **20**. For Hs-FIGL, Dm-spastin, Mm-VCP/p97, Xl-katanin, and Hs-PCH2, compound **19** (1 mM in DMSO) was diluted 1:25 in buffer to achieve 2X compound. For typical conditions, 2X compound (10 μL) was added to protein/NADH mix (8 μL) and incubated for 10 min at RT. The reaction was initiated with the addition of 10 mM MgATP (2 μL), and the fluorescence signal was monitored using a Synergy NEO Microplate Reader (λex = 340 nm, 440 nm emission filter). The slope of the fluorescence values versus time was calculated to give the rate of fluorescence decrease. The ATPase rate was calculated from the rate of fluorescence decrease from a standard curve measuring the contribution of ADP to the NADH fluorescence signal.

Assay buffers:

GFP-Sc-Dyn and its mutants (10-45 nM): 30 mM HEPES pH 7.4, 50 mM KOAc, 1 mM EGTA, 2 mM MgOAc, 1 mM DTT, 0.01% Triton™ X-100

Sc-Dyn-lyso (5 nM): 30 mM HEPES pH 7.4, 50 mM KOAc, 1 mM EGTA, 2 mM MgOAc, 1 mM DTT, 10% glycerol, 0.01% Triton™ X-100

Xl-katanin (50 nM): 25 mM K-HEPES pH 7.5, 70 mM KCl, 20 mM (NH4)_2_SO_4_, 5 mM MgCl_2_, 2.5 mM DTT, 0.01% w/v Triton™ X-100

Mm-p97 (400 nM): 30 mM HEPES pH 7.4, 50 mM KOAc, 1 mM EGTA, 2 mM MgOAc, 1 mM DTT, 0.01% w/v Triton™ X-100

Hs-FIGL1 (50 nM): 25 mM Na-MES pH 6.5, 70 mM KOAc, 20 mM (NH4)_2_SO_4_, 5 mM Mg(OAc)_2_, 1 mM TCEP, 0.01% w/v Triton™ X-100

Dm-spastin (75 nM): 25 mM K-HEPES pH 7.5, 225 mM KCl, 2.5 mM (NH4)_2_SO_4_, 5 mM MgCl_2_, 2.5 mM DTT, 1 mg/mL BSA, 0.005% w/v Triton™ X-100

### Microscale thermophoresis assay (MST)

The binding affinities of compound **20** to Sc-Dyn-lysoMut was measured using Monolith NT.115 (NanoTemper Technologies, Munich, Germany). Sc-Dyn-lysoMut was labeled with the Monolith NT Protein Labeling Kit RED-NHS. To obtain a fluorescence intensity between 200 and 1100 counts at 20% LED power, a final concentration of 30-50 nM protein was used. The MST assays were performed in the following buffer: 20 mM Tris-HCl pH 8.0, 50 mM K-Ac, 2 mM Mg(Ac)_2_, 1 mM EGTA, 10% glycerol, 0.005% Triton-X100. A 50X compound stock was prepared by a 1:2 serial dilution of the compound in DMSO starting from 20 mM (12 doses total). The 50X compound stock was diluted in buffer to obtain a 2X compound stock. The 2X protein stock was mixed well with 2X compound and incubated for 10 min at RT in the dark. Each sample was transferred to a Monolith premium capillary tube, and measurements were performed at at 20% and 40% MST power for Sc-Dyn-lyso and GFP-Sc-Dyn constructs, respectively. A binding curve was observed at 1.5s after the start of thermophoresis. The F_norm_ values were fitted to Equation (2) using MATLAB (Natick, Ma) at different concentrations of compound **20**.

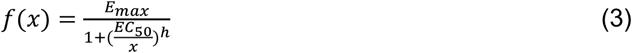

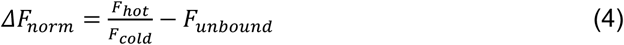

In Equation (3), *E_max_* is the maximum response, EC_50_ is the compound concentration that produces the half-maximum response, and h corresponds to the hill slope. In Equation (4) *F_hot_* denotes the fluorescence value at the hot state and *F_hot_* represents the fluorescence value at the cold state. *F_unbound_* indicates the mean fluorescence at the unbound state.

### Single-molecule motility assays

Single-molecule TIRF assays were carried out using a Nikon Eclipse Ti inverted microscopy with a N1-1.49 100x Plan Apo objective (Nikon). The microscope was outfitted with 488 nm (Spectra-physics Cyan Scientific 40 mW) and 561 nm (Cobalt Jive 50 mW) lasers, dichroic mirrors (Semrock Di01-R488 and Di01-R488/561), excitation filters (Semrock FF01-488/6 and FF01-482/563), and emission filters (FF01-525/45 and FF01-609/54). Images were captured using a Photometric Prime 95B sCMOS camera. Flow cells were constructed using glass slides (Ted Pella 260600), glass coverslips (Fisher 12-541A), and double-sided tape. Cover slips were thoroughly washed with 70% ethanol and water prior to use. All single-molecule experiments were carried out at 23° C.

Rhodamine-labeled microtubules were polymerized from bovine tubulin (~10% final rhodamine-labeled tubulin, prepared as previously and stabilized with 10 μM taxol (Sigma T7191) before being pelleted in a TLA120.1 rotor at 90,000 rpm for 10 min at 25°C (Beckman Coulter) (Subramanian et al., 2013; Ti et al., 2016). The pellet was washed and resuspended with BRB80 buffer plus 10 μM taxol. Microtubules were attached to the κ-casein-blocked coverslip surface using rigor-kinesin as previously described (Mickolajczyk et al., 2016).

GFP-labeled dynein GST dimers, prepared as previously, were first incubated in dynein motility buffer (30 mM HEPES pH=7.2, 50 mM K-Acetate, 2 mM Mg-Acetate, 1 mM EGTA, 10% v/v glycerol) containing 50 μM compound **20** in 1% DMSO and incubated on ice for 30 minutes (Niekamp et al., 2019). Motors were drawn from this stock and diluted 10,000-fold (final concentration 100 pM total dynein) into motility buffer containing 0-50 μM compound **20** in 1% DMSO as well as 100 μM MgATP. In the final serial dilution, the motility buffer was supplemented with 1 mM DTT, 1 mM Mg-ATP, 0.05 mg/mL κ-casein, 10 μM taxol, 20 mM glucose, 20 μM/mL glucose oxidase, and 8 μg/mL catalase. The final mixture was added to flow cells containing immobilized microtubules. A single image of rhodamine microtubules was taken first, then a movie of GFP-dynein was taken in the same field of view at 5 frames per second. Landing rates were determined by manual kymograph analysis in ImageJ (NIH). Landing rate data were fitted with a sigmoidal dose response equation:

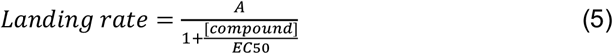

Where A gives a Y-intercept corresponding to the landing rate in the absence of drug, and IC_50_ gives the drug concentration yielding A/2. The run lengths and velocities of individual dyneins were determined by linear fitting to particle trajectories obtained using FIESTA software (Ruhnow et al., 2011). Population velocities were obtained by taking the sample mean, and population run lengths were obtained by fitting the cumulative density function to an offset exponential (Thompson et al., 2013; Thorn et al., 2000). All data analysis was performed in MATLAB (Mathworks).

### Chemical Synthesis

General. Solvents and reagents were purchased from VWR or Sigma Aldrich. All reactions involving air- or moisture-sensitive compounds were performed under nitrogen atmosphere using dried glassware. For compounds **3-19**, ^1^H and ^13^C (for **6** and **19**) NMR spectra were recorded at 500MHz and 125MHz respectively, on a Bruker Advance III HD 500 MHz NMR spectrometer equipped with at TCI cryogenic probe with enhanced ^1^H and ^13^C detection. For compound **20**, ^1^H and ^13^C NMR spectra were recorded at 600MHz and 151M Hz respectively, on a Bruker Advance II 600 MHz NMR spectrometer eqiupped with a 5 mm TXI cryogenic probe. All data was collected at 298K, signals were reported in ppm, internally referenced for ^1^H and ^13^C to chloroform signal at 7.26 ppm or 77.0 ppm; to DMSO signal at 2.50 ppm or 39.5 ppm, or TMS at 0 ppm. Chemical shifts are reported in parts per million (ppm) and the coupling constants (*J*) are expressed in hertz (Hz). Splitting patterns are designated as follows: s, singlet; d, doublet; t, triplet; m, multiplet; dd, doublet of doublets; ddd, double of doublets of doublets; dt, doublet of triplets. Flash chromatography purifications were performed on Combi*Flash* Rf (Teledyne ISCO) as the stationary phase. Purity for all tested compounds was determined through high-performance liquid chromatography and all compounds were found to be >95% pure. Liquid chromatography mass spectral analyses were obtained using a Waters Acquity H-Class UPLC/MS with a QDa mass spectrometer. The system used a eλ Photodiode Array Detector detector, and a Symmetry C18 (3.5 micron) 2.1 x 50 mm column for separation (mobile phase for positive mode: solvent A: water with 0.1% formic acid, solvent B: acetonitrile with 0.1% formic acid). Values are reported in units of mass to charge (m/z).

**General method A** (used to prepare compounds **6, 9, 10, 13, 17, 18, 19, 20**)

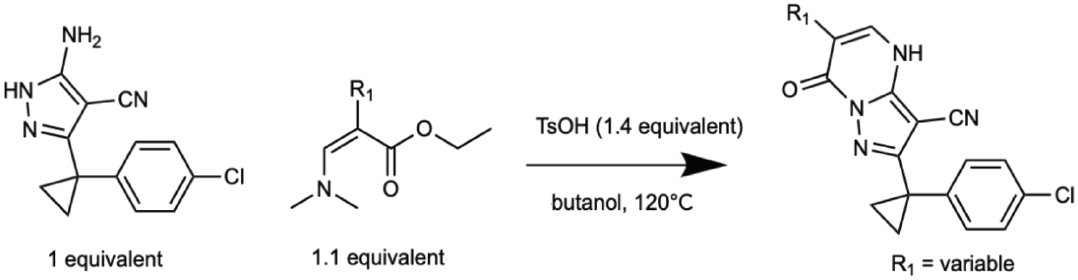

A mixture of the appropriate ethyl or methyl (Z)-2-(alkyloxy)-3-(dimethylamino)prop-2-enoate (1.1 equivalents), 5-amino-3-[1-(4-chlorophenyl)cyclopropyl]-1H-pyrazole-4-carbonitrile (1 equivalent), and *p*-toluenesulfonic acid hydrate (1.4 equivalents) in butanol was stirred at 120 °C for 16 h. The reaction mixture was concentrated *in vacuo*. The residue was purified column chromatography (silica gel, hexane/ethyl acetate) and washed with ethyl acetate to give the desired product.

**General method B** (used to prepare compound **16**)

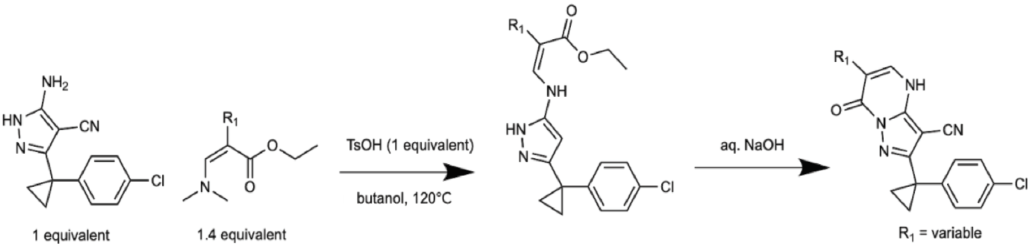

To a solution of ethyl (E)-3-(dimethylamino)-2-(aryloxy)prop-2-enoate (1.4 equivalents) in butanol were added 5-amino-3-[1-(4-chlorophenyl) cyclopropyl]-1H-pyrazole-4-carbonitrile (1 equivalent), *p*-toluenesulfonic acid hydrate (1 equivalent). The mixture was stirred at 120 °C for 3 days. The mixture was cooled to room temperature and concentrated *in vacuo*. The residue was purified by column chromatography (0% - 60% hexane in ethyl acetate) to give the corresponding ethyl (E)-3-[[3-[1-(4-chlorophenyl)cyclopropyl]-4-cyano-1H-pyrazol-5-yl]amino]-2-(aryloxy)prop-2-enoate, which was subsequently dissolved in ethanol and treated with aqueous sodium hydroxide (1.8 equivalents NaOH) at room temperature. The mixture was concentrated after being stirred at room temperature for 6 h. The mixture was neutralized with 1N HCl and extracted with ethyl acetate. The combined organic layer was washed with water and brine, dried over MgSO_4_, filtered and concentrated *in vacuo*. The residue was purified by column chromatography (silica-gel, 50% - 100% hexane in ethyl acetate) to give the desired product.

**General method C** (used to prepare compound **4**)

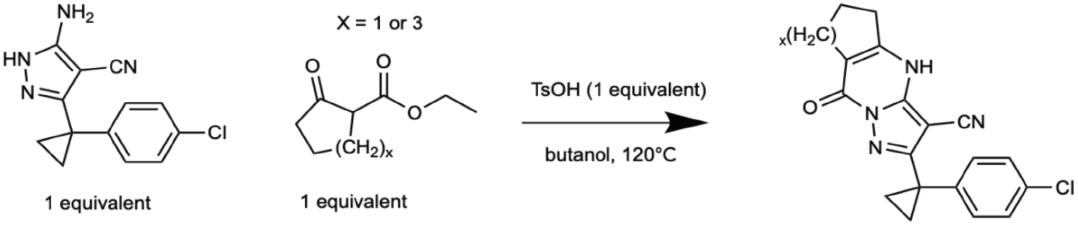

A mixture of the appropriate ethyl 2-oxocycloalkane carboxylate (1 equivalent), 5-amino-3-[1-(4-chlorophenyl)cyclopropyl]-1H-pyrazole-4-carbonitrile (1 equivalent) and *p*-toluenesulfonic acid hydrate (1 equivalent) in butanol was stirred at 120 °C for 1.5 h. The reaction was concentrated *in vacuo*. The residue was purified by column chromatography (silica-gel, 50-100% ethyl acetate in hexane) to give the desired product.

**General method D** (used to prepare **7**)

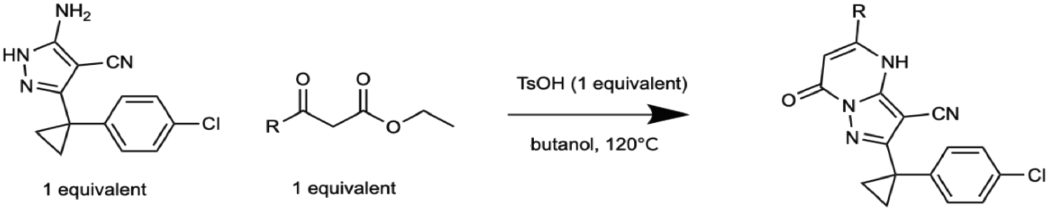

A mixture of 5-amino-3-[1-(4-chlorophenyl)cyclopropyl]-1H-pyrazole-4-carbonitrile (1 equivalent), the appropriate 3-substituted ethyl-3-oxopropanoate (1 equivalent) and *p*-toluenesulfonic acid hydrate (1 equivalent) in butanol was stirred at 120°C for 2 h. The reaction was cooled to room temperature. Resulting precipitates were collected and washed with methanol to give the desired product.

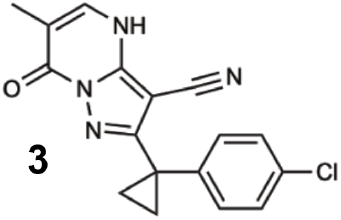

To a solution of methyl (E)-3-(dimethylamino)-2-methyl-prop-2-enoate (180 mg, 1.26 mmol) in butanol were added *p*-toluenesulfonic acid hydrate (359 mg, 1.89 mmol) and 3-amino-5-[1-(4-chlorophenyl)cyclopropyl]-1H-pyrazole-4-carbonitrile (326 mg, 1.26 mmol). The mixture was stirred at 120 °C for 16 h. The mixture was poured into water and extracted with EtOAc. The organic layer was washed with water, then washed with brine, dried over magnesium sulfate, and concentrated in vacuo. The residue was purified by column chromatography (silica-gel, 10%-60% ethyl acetate in hexane) to give a crude product (140 mg). 29 mg of the crude material was purified with preparative HPLC (water/CH3CN, 0.1% formic acid) to give 2-[1-(4-chlorophenyl) cyclopropyl]-6-methyl-7-oxo-4H-pyrazolo[1,5-a]pyrimidine-3-carbonitrile (**3**, 16 mg, 49 μmol, 3.91% yield) as a colorless solid. 1H NMR (500 MHz, DMSO-d6) δ 13.22 (s, 0H), 7.71 (s, 1H), 7.47 – 7.14 (m, 4H), 1.94 (s, 3H), 1.46 (s, 2H), 1.30 (s, 2H). MS m/z 325.0 [M+1]+.

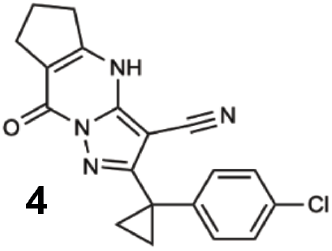

A mixture of ethyl 2-oxocyclopentanecarboxylate (30.2 mg, 193 μmol, 27.9 uL), 3-amino-5-[1-(4-chlorophenyl)cyclopropyl]-1H-pyrazole-4-carbonitrile (50.0 mg, 193 μmol) and *p*-toluenesulfonic acid hydrate (36.8 mg, 193 μmol) in butanol (1.50 mL) was stirred at 120 °C for 1.5 h. The reaction was concentrated in vacuo. The residue was purified by column chromatgraphy (silica-gel, 50-100% EtOAc in hexane) to give 2-(1-(4-chlorophenyl)cyclopropyl)-8-oxo-5,6,7,8-tetrahydro-4H-cyclopenta[d]pyrazolo[1,5-a]pyrimidine-3-carbonitrile (**4**, 44.7 mg, 127 μmol, 66% yield) as an off white solid. 1H NMR (500 MHz, DMSO-d6) δ 13.50 (s, 1H), 7.36 (d, J = 7.9 Hz, 2H), 7.28 (d, J = 7.8 Hz, 2H), 2.91 (t, J = 7.6 Hz, 2H), 2.69 (t, J = 7.2 Hz, 2H), 2.13 – 2.03 (m, 2H), 1.50 (s, 2H), 1.37 (s, 2H). MS m/z: 351.2 [M+1]+.

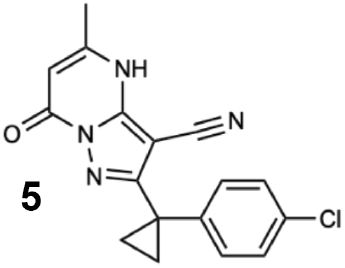

The mixture of ethyl 3-oxobutanoate (50.0 mg, 384 μmol), 3-amino-5-[1-(4-chlorophenyl)cyclopropyl]-1H-pyrazole-4-carbonitrile (99.4 mg, 384 μmol) and *p*-toluenesulfonic acid hydrate (73.1 mg, 384 μmol) in BuOH (3.0 mL) was stirred at 120 °C for 2 h. The reaction mixture was cooled to room temperature. The precipitate was collected and washed with MeOH to give 2-[1-(4-chlorophenyl) cyclopropyl]-5-methyl-7-oxo-4H-pyrazolo[1,5-a] pyrimidine-3-carbonitrile (**5**, 62.4 mg, 192 μmol, 50% yield) as an off-white solid. ^1^H NMR (500 MHz, DMSO-*d*_6_) δ 13.17 (s, 1H), 7.38 – 7.34 (m, 2H), 7.31 – 7.26 (m, 2H), 5.83 (s, 1H), 2.29 (s, 3H), 1.50 (s, 2H), 1.37 (s, 2H). MS m/z: 325.1 [M+1]+.

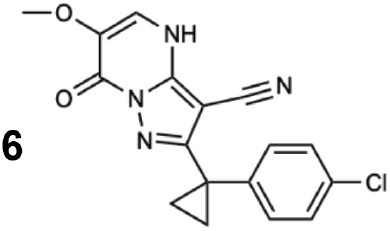

General method A was used to obtain 2-[1-(4-chlorophenyl)cyclopropyl]-6-methoxy-7-oxo-4H-pyrazolo [1,5-a]pyrimidine-3-carbonitrile (**6**, 66% yield). 1H NMR (500 MHz, DMSO-d6) δ 13.18 (s, 1H), 7.77 (s, 1H), 7.38 (d, J= 8.5 Hz, 2H), 7.31 (d, J= 8.3 Hz, 2H), 3.77 (s, 3H), 1.53 (q, J= 4.5 Hz, 2H), 1.39 (q, J= 4.5 Hz, 2H). ^13^C NMR (151 MHz, DMSO-*d*_6_) δ 160.50, 152.85, 145.34, 141.15, 135.74, 131.76, 131.76, 130.24, 130.24, 128.79, 123.74, 112.92, 73.87, 58.79, 23.95, 15.90, 15.90 MS m/z: 341.1 [M+1]+.

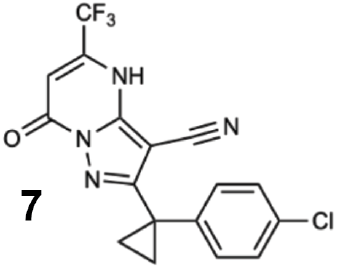

General method D was used to obtain 2-(1-(4-chlorophenyl)cyclopropyl)-7-oxo-5-(trifluoromethyl)-4,7-dihydropyrazolo[1,5-a]pyrimidine-3-carbonitrile (**7**, 7.7% yield)1H NMR (500 MHz, DMSO-d6) δ 7.34 (d, J = 7.9 Hz, 2H), 7.26 (d, J = 7.6 Hz, 2H), 5.92 (s, 1H), 1.49 (s, 2H), 1.32 (s, 2H). 1H was hidden. MS; m/z 379.1 (M+1).

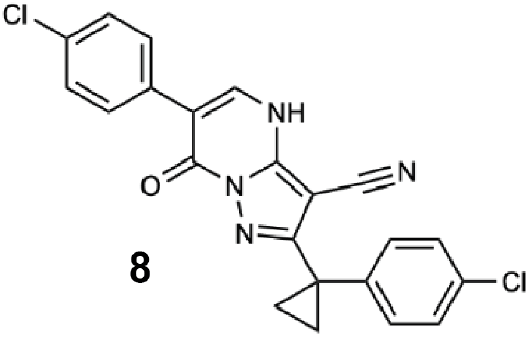

To a solution of methyl (Z)-2-(4-chlorophenyl)-3-(dimethylamino)prop-2-enoate (111 mg, 464 μmol) in butanol (5.00 mL) were added 1 (100 mg, 387 μmol), *p*-toluenesulfonic acid hydrate (74 mg, 387 μmol). The mixture was stirred at 120 °C for 3 hours. The mixture was cooled to room temperature. The resulting solid was collected by filtration using methanol to give 6-(4-chlorophenyl)-2-[1-(4-chlorophenyl)cyclopropyl]-7-oxo-4H-pyrazolo[1,5-a]pyrimidine-3-carbonitrile (**8**, 74 mg, 176 μmol, 45% yield) as a white solid. 1H NMR (500 MHz, DMSO-d6) δ 13.88 (s, 1H), 8.19 (s, 1H), 7.76 – 7.68 (m, 2H), 7.56 – 7.48 (m, 2H), 7.44 – 7.37 (m, 2H), 7.36 – 7.30 (m, 2H), 1.60 – 1.52 (m, 2H), 1.46 – 1.39 (m, 2H). MS m/z: 419 [M-H]-.

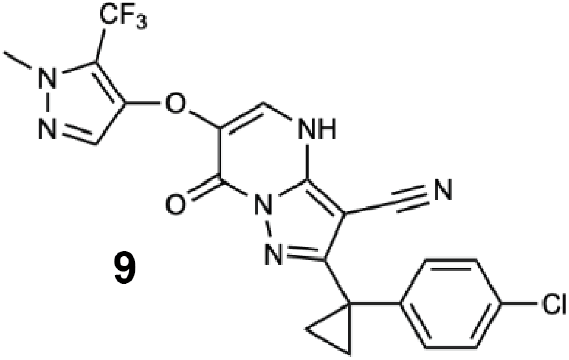

General method A was used to obtain 2-[1-(4-chlorophenyl)cyclopropyl]-6-[1-methyl-5-(trifluoromethyl) pyrazol-3-yl]oxy-7-oxo-4H-pyrazolo[1,5-a]pyrimidine-3-carbonitrile (**9**, 43% yield). 1H NMR (500 MHz, DMSO-d6) δ 8.00 (s, 1H), 7.44 – 7.36 (m, 2H), 7.35 – 7.27 (m, 2H), 6.34 (s, 1H), 3.80 (s, 3H), 1.55 – 1.47 (m, 2H), 1.38 – 1.30 (m, 2H). MS m/z: 475.2 [M+1]+.

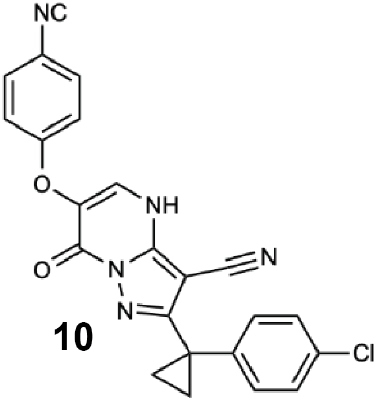

General method A was used to obtain 2-[1-(4-chlorophenyl)cyclopropyl]-6-(4-cyanophenoxy)-7-oxo-4H-pyrazolo[1,5-a]pyrimidine-3-carbonitrile (**10**, 9.9% yield). 1H NMR (500 MHz, DMSO-d6) δ 7.95 (s, 1H), 7.76 (d, J = 8.9 Hz, 2H), 7.41 – 7.35 (m, 2H), 7.34 – 7.27 (m, 2H), 7.06 (d, J = 8.9 Hz, 2H), 1.54 – 1.48 (m, 2H), 1.35 – 1.30 (m, 2H). MS m/z: 428.2 [M+1]+.

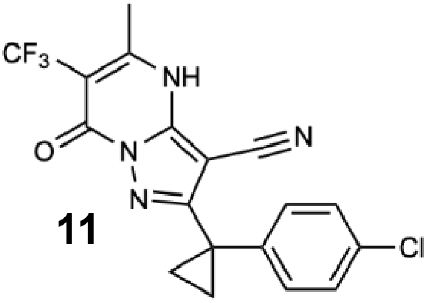

A mixture of 5-amino-3-[1-(4-chlorophenyl)cyclopropyl]-1H-pyrazole-4-carbonitrile (100 mg, 387 μmol), ethyl 3-oxo-2-(trifluoromethyl)butanoate (77 mg, 387 μmol), and *p*-toluenesulfonic acid hydrate (74 mg, 387 μmol) in butanol (5.00 mL) was stirred at 100 °C for 8 h. The reaction was concentrated *in vacuo*. The residue was purified by column chromatography (silica-gel, 20-80% ethyl acetate in hexane) and washed with diisopropyl ether to give 2-[1-(4-chlorophenyl)cyclopropyl]-5-methyl-7-oxo-6-(trifluoromethyl)-4H-pyrazolo[1,5-a]pyrimidine-3-carbonitrile (**11**, 56.0 mg, 143 μmol, 37% yield) as a pale yellow solid. ^1^H NMR (500 MHz, Chloroform-*d*) δ 10.99 (s, 1H), 7.32 (d, *J* = 8.4 Hz, 2H), 7.28 – 7.23 (m, 2H), 2.53 – 2.48 (m, 3H), 1.71 – 1.66 (m, 2H), 1.36 (q, *J* = 4.5 Hz, 2H). MS m/z 393.318 [M+1]^+^.

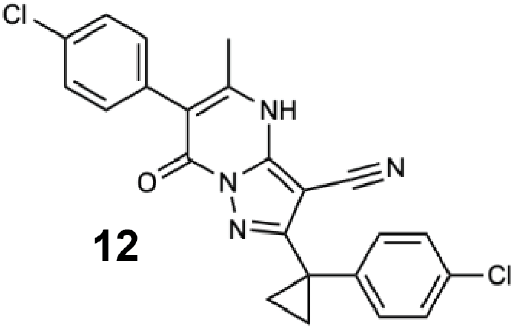

A mixture of 2-[1-(4-chlorophenyl)cyclopropyl]-6-iodo-5-methyl-7-oxo-4H,7H-pyrazolo [1,5-a]pyrimidine-3-carbonitrile (**14**, 10 mg, 22 μmol),cyclopentyl(diphenyl)phosphane; dichloromethane;dichloropalladium;iron (1.83 mg, 2.22 μmol), potassium carbonate (6.1 mg, 44.4 μmol), and (4-chlorophenyl)boronic acid (4.2 mg, 26.7 μmol) in EtOH (0.5 mL) and toluene (0.5 mL) was stirred at 80 °C for 5.5 h. The reaction was concentrated in vacuo. The residue was purified by column chromatography (silica-gel, 0-100% ethyl acetate in hexane) to give 6-(4-chlorophenyl)-2-[1-(4-chlorophenyl)cyclopropyl]-5-methyl-7-oxo-4H-pyrazolo[1,5-a]pyrimidine-3-carbonitrile (**12**, 3.6 mg, 8.3 μmol, 37% yield) as a yellow solid. 1H NMR (500 MHz, DMSO-d6) δ 13.30 (s, 1H), 7.50 (d, J = 8.0 Hz, 2H), 7.37 (d, J = 8.5 Hz, 2H), 7.34 – 7.26 (m, 4H), 2.16 (s, 3H), 1.39 (s, 2H), 1.23 (s, 2H). MS m/z 435.3 [M+1]+.

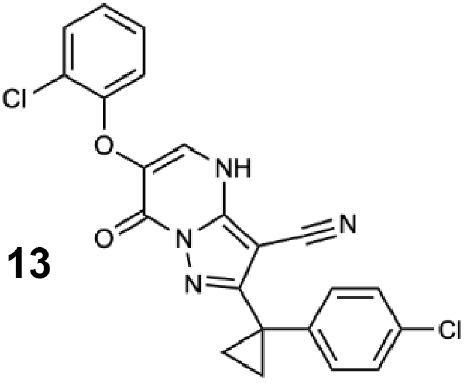

General method A was used to obtain 6-(2-chlorophenoxy)-2-[1-(4-chlorophenyl) cyclopropyl]-7-oxo-4H-pyrazolo[1,5-a]pyrimidine-3-carbonitrile (**13**, 20% yield). 1H NMR (500 MHz, Chloroform-d) δ 8.09 (s, 1H), 7.48 (d, J = 7.9 Hz, 1H), 7.36 (d, J = 8.6 Hz, 2H), 7.30 (d, J = 8.6 Hz, 2H), 7.22 – 7.15 (m, 1H), 7.01 (t, J = 8.3 Hz, 1H), 6.94 – 6.86 (m, 1H), 1.53 – 1.47 (m, 2H), 1.35 (t, J = 5.6 Hz, 2H). MS m/z: 437.2 [M+1]+.

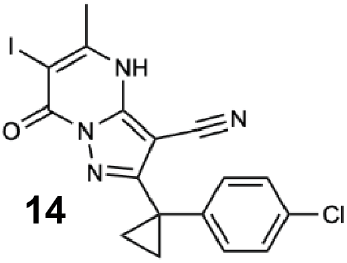

A mixture of ethyl 3-oxobutanoate (50.0 mg, 384 μmol, 48.6 uL), 3-amino-5-[1-(4-chlorophenyl)cyclopropyl]-1H-pyrazole-4-carbonitrile (99.4 mg, 384 μmol) and *p*-toluenesulfonic acid hydrate (73.1 mg, 384 μmol) in butanol (3.0 mL) was stirred at 120 °C for 2 h. The reaction was cooled to room temperature. The precipitate was collected and washed with MeOH to give 2-[1-(4-chlorophenyl)cyclopropyl]-6-iodo-5-methyl-7-oxo-4H,7H-pyrazolo [1,5-a]pyrimidine-3-carbonitrile (**14**, 62.4 mg, 192 μmol, 50% yield) as an off white solid. 1H NMR (500 MHz, DMSO-d6) δ 13.17 (s, 1H), 7.38 – 7.34 (m, 2H), 7.31 – 7.26 (m, 2H), 5.83 (s, 1H), 2.29 (s, 3H), 1.50 (s, 2H), 1.37 (s, 2H). MS m/z 325.1 [M+1]+.

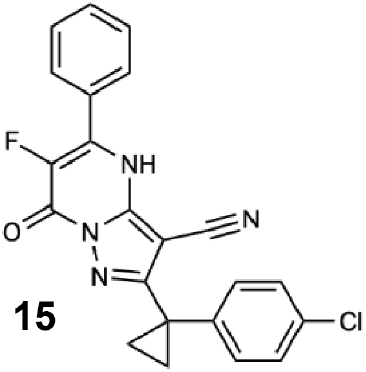

A mixture of 1 (54.5 mg, 211 μmol), ethyl (E)-3-(dimethylamino)-2-fluoro-3-phenyl-prop-2-enoate (50.0 mg, 211 μmol) and *p*-toluenesulfonic acid hydrate (43.9 mg, 255 mol) in butanol (2.0 mL) was stirred at 130 °C for 2 h. The reaction mixture was concentrated in vacuo. The residue was purified by column chromatography (silica-gel, 50-80%, ethyl acetate in hexane). The resulting solid was triturated with ethyl acetate to give 2-[1-(4-chlorophenyl)cyclopropyl]-6-fluoro-7-oxo-5-phenyl-4H-pyrazolo[1,5-a] pyrimidine-3-carbonitrile (**15**, 29 mg, 72.1 μmol, 34% yield) as a colorless solid. 1H NMR (500 MHz, DMSO-d6) δ 7.82 – 7.65 (m, 2H), 7.66 – 7.46 (m, 3H), 7.37 (d, J= 8.1 Hz, 2H), 7.30 (d, J= 8.2 Hz, 2H), 1.62 – 1.46 (m, 2H), 1.45 – 1.28 (m, 2H). MS m/z 405.1 [M+1]+.

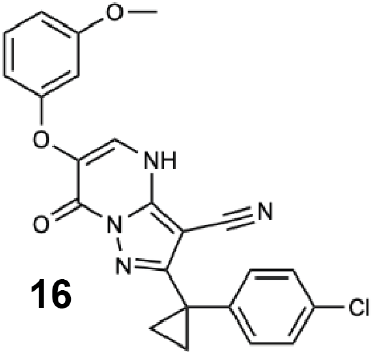

General method B was used to obtain 2-[1-(4-chlorophenyl)cyclopropyl]-6-(3-methoxyphenoxy)-7-oxo-4H-pyrazolo[1,5-a]pyrimidine-3-carbonitrile (**16**, 80% yield). ^1^H NMR (500 MHz, DMSO-d6) δ 13.68 (s, 1H), 8.21 (s, 1H), 7.38 (d, J = 8.5 Hz, 2H), 7.32 (d, J = 8.5 Hz, 2H), 7.21 – 7.16 (m, 1H), 6.65 – 6.57 (m, 3H), 3.73 (s, 3H), 1.56 – 1.49 (m, 2H), 1.43 – 1.33 (m, 2H). MS m/z: 433.1 [M+1]^+^.

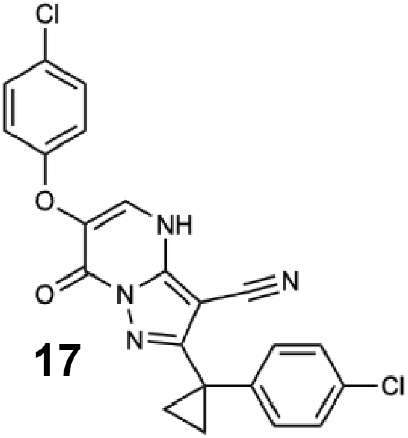

General method A was used to obtain 6-(4-chlorophenoxy)-2-(1-(4-chlorophenyl) cyclopropyl)-7-oxo-4,7-dihydropyrazolo[1,5-*a*]pyrimidine-3-carbonitrile (**17**, 14.08% yield). 1H NMR (500 MHz, DMSO-d6) δ 13.69 (s, 1H), 8.09 (s, 1H), 7.48 – 7.24 (m, 6H), 7.00 (d, J = 6.9 Hz, 2H), 1.50 (s, 2H), 1.35 (s, 2H). MS m/z: 437.1 [M+1]+.

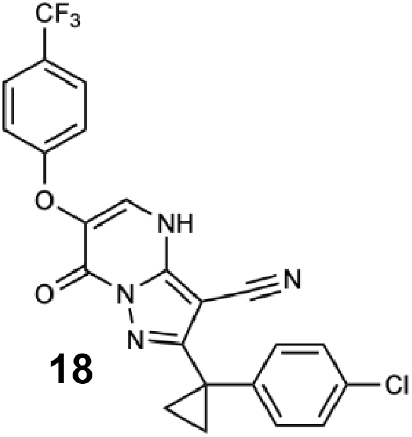

General method A was used to obtain 2-[1-(4-chlorophenyl)cyclopropyl]-7-oxo-6-[4-(trifluoromethyl) phenoxy]-4H-pyrazolo[1,5-a]pyrimidine-3-carbonitrile (**18**, 17% yield) as a pale yellow solid. 1H NMR (500 MHz, DMSO-d6) δ 8.15 (s, 1H), 7.66 (d, J = 8.4 Hz, 2H), 7.49 – 7.04 (m, 6H), 1.64 – 1.28 (m, 4H). MS m/z: 469 [M-H]-.

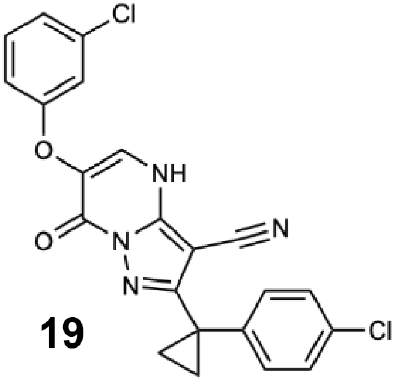

General method A was used to obtain 6-(3-chlorophenoxy)-2-[1-(4-chlorophenyl) cyclopropyl]-7-oxo-4H-pyrazolo[1,5-a]pyrimidine-3-carbonitrile (**19**, 10% yield) as a beige solid. ^1^H NMR (500 MHz, DMSO-*d*_6_) δ 8.26 (s, 1H), 7.38 (d, *J*= 8.6 Hz, 2H), 7.36 – 7.29 (m, 3H), 7.18 (d, *J*= 2.3 Hz, 1H), 7.10 (dd, *J*= 8.0, 1.9 Hz, 1H), 7.06 (dd, *J*= 8.4, 2.5 Hz, 1H), 1.57 – 1.46 (m, 2H), 1.45 – 1.34 (m, 2H). ^13^C NMR (151 MHz, DMSO-*d*_6_) δ 163.49, 160.09, 159.87, 153.04, 141.95, 134.10, 131.48, 131.31, 130.13, 130.13, 128.69, 128.69, 128.24, 122.09, 115.46, 115.46, 114.68, 114.68, 75.88, 24.15, 15.73, 15.73. MS m/z: 439 [M+H]^+^.

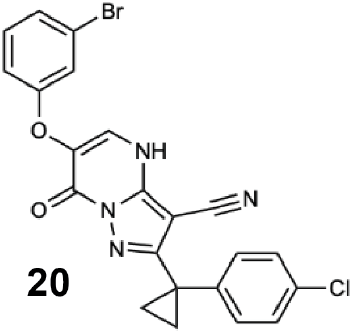

General method A was used to obtain 6-(3-bromophenoxy)-2-[1-(4-chlorophenyl) cyclopropyl]-7-oxo-4H-pyrazolo[1,5-a]pyrimidine-3-carbonitrile (**20**, 67% yield) as a beige solid. ^1^H NMR (600 MHz, DMSO-rá) δ 8.26 (s, 1H), 7.38 (d, *J* = 8.4 Hz, 2H), 7.35 – 7.29 (m, 3H), 7.29 – 7.21 (m, 2H), 7.10 (d, *J* = 7.9 Hz, 1H), 1.52 (d, *J* = 5.8 Hz, 2H), 1.40 (d, *J* = 4.7 Hz, 2H). ^13^C NMR (151 MHz, DMSO-*d*_6_) δ 159.84, 159.01, 152.47, 146.43, 140.70, 134.24, 131.34, 131.23, 129.83, 128.33, 128.21, 125.19, 122.09, 117.84, 114.87, 112.37, 74.96, 23.53, 15.37. MS m/z: 479.2 [M-H]^-^.

### Data availability statement

Cryo-EM maps have been deposited in the EMDB under accession codes EMD-XXXX (Map 1), EMD-XXXX (Map 2), EMD-XXXX (Map 3), EMD-XXXX (Map 4), EMD-XXXX (Map 5). Coordinates are available from the RCSB Protein Data Bank under accession codes XXXX (Map 1), XXXX (Map 2), XXXX (Map 3), XXXX (Map 4), and XXXX (Map 5). The crystal structure of Sc-DynlysoMut has been deposited in RCSB Protein Data Bank under accession code XXXX. All other data generated or analyzed during this study are available in this article or from the corresponding author on reasonable request.

## References

Afonine, P.V., Klaholz, B.P., Moriarty, N.W., Poon, B.K., Sobolev, O.V., Terwilliger, T.C., Adams, P.D., and Urzhumtsev, A. (2018). New tools for the analysis and validation of cryo-EM maps and atomic models. Acta Crystallogr D Struct Biol 74, 814–840.

Arkhipova, V., Guskov, A., and Slotboom, D.-J. (2017). Analysis of the quality of crystallographic data and the limitations of structural models. J. Gen. Physiol. 149, 1091–1103.

Banerjee, S., Bartesaghi, A., Merk, A., Rao, P., Bulfer, S.L., Yan, Y., Green, N., Mroczkowski, B., Neitz, R.J., Wipf, P., et al. (2016). 2.3 Å resolution cryo-EM structure of human p97 and mechanism of allosteric inhibition. Science 351, 871–875.

Bhabha, G., Cheng, H.-C., Zhang, N., Moeller, A., Liao, M., Speir, J.A., Cheng, Y., and Vale, R.D. (2014). Allosteric communication in the dynein motor domain. Cell 159, 857–868.

Burgess, S.A., Walker, M.L., Sakakibara, H., Knight, P.J., and Oiwa, K. (2003). Dynein structure and power stroke. Nature 421, 715–718.

Carter, A.P., Cho, C., Jin, L., and Vale, R.D. (2011). Crystal structure of the dynein motor domain. Science 331, 1159–1165.

Cho, C., Reck-Peterson, S.L., and Vale, R.D. (2008). Regulatory ATPase sites of cytoplasmic dynein affect processivity and force generation. J. Biol. Chem. 283, 25839–25845.

Cupido, T., Pisa, R., Kelley, M.E., and Kapoor, T.M. (2019). Designing a chemical inhibitor for the AAA protein spastin using active site mutations. Nat. Chem. Biol. 15, 444–452.

DeWitt, M.A., Cypranowska, C.A., Cleary, F.B., Belyy, V., and Yildiz, A. (2015). The AAA3 domain of cytoplasmic dynein acts as a switch to facilitate microtubule release. Nat. Struct. Mol. Biol. 22, 73–80.

Dharan, A., and Campbell, E.M. (2018). Role of Microtubules and Microtubule-Associated Proteins in HIV-1 Infection. J. Virol. 92.

Elshenawy, M.M., Canty, J.T., Oster, L., Ferro, L.S., Zhou, Z., Blanchard, S.C., and Yildiz, A. (2019). Cargo adaptors regulate stepping and force generation of mammalian dynein–dynactin. Nature Chemical Biology 15, 1093–1101.

Emsley, P., Lohkamp, B., Scott, W.G., and Cowtan, K. (2010). Features and development of Coot. Acta Crystallogr. D Biol. Crystallogr. 66, 486–501.

Erzberger, J.P., and Berger, J.M. (2006). Evolutionary relationships and structural mechanisms of AAA+ proteins. Annu. Rev. Biophys. Biomol. Struct. 35, 93–114.

Firestone, A.J., Weinger, J.S., Maldonado, M., Barlan, K., Langston, L.D., O’Donnell, M., Gelfand, V.I., Kapoor, T.M., and Chen, J.K. (2012). Small-molecule inhibitors of the AAA+ ATPase motor cytoplasmic dynein. Nature 484, 125–129.

Gibbons, B.H., and Gibbons, I.R. (1987). Vanadate-sensitized cleavage of dynein heavy chains by 365-nm irradiation of demembranated sperm flagella and its effect on the flagellar motility. J. Biol. Chem. 262, 8354–8359.

Höing, S., Yeh, T.-Y., Baumann, M., Martinez, N.E., Habenberger, P., Kremer, L., Drexler, H.C.A., Küchler, P., Reinhardt, P., Choidas, A., et al. (2018). Dynarrestin, a Novel Inhibitor of Cytoplasmic Dynein. Cell Chemical Biology 25, 357–369.e6.

Jerabek-Willemsen, M., André, T., Wanner, R., Roth, H.M., Duhr, S., Baaske, P., and Breitsprecher, D. (2014). MicroScale Thermophoresis: Interaction analysis and beyond. J. Mol. Struct. 1077, 101–113.

Johnson, R.L., Rothman, A.L., Xie, J., Goodrich, L.V., Bare, J.W., Bonifas, J.M., Quinn, A.G., Myers, R.M., Cox, D.R., Epstein, E.H., Jr, et al. (1996). Human homolog of patched, a candidate gene for the basal cell nevus syndrome. Science 272, 1668–1671.

Kabsch, W. (2010). XDS. Acta Crystallogr. D Biol. Crystallogr. 66, 125–132.

Kon, T., Nishiura, M., Ohkura, R., Toyoshima, Y.Y., and Sutoh, K. (2004). Distinct functions of nucleotide-binding/hydrolysis sites in the four AAA modules of cytoplasmic dynein. Biochemistry 43, 11266–11274.

Kon, T., Oyama, T., Shimo-Kon, R., Imamula, K., Shima, T., Sutoh, K., and Kurisu, G. (2012). The 2.8 Å crystal structure of the dynein motor domain. Nature 484, 345–350.

McCoy, A.J., Grosse-Kunstleve, R.W., Adams, P.D., Winn, M.D., Storoni, L.C., and Read, R.J. (2007). Phaser crystallographic software. J. Appl. Crystallogr. 40, 658–674.

Mickolajczyk, K.J., Deffenbaugh, N.C., Ortega-Arroyo, J., Andrecka, J., Kukura, P., and Hancock, W.O. (2016). Kinetics of Nucleotide-Dependent Structural Transitions in the Kinesin-1 Hydrolysis Cycle. Biophysical Journal 110, 193a.

Mijalkovic, J., Prevo, B., Oswald, F., Mangeol, P., and Peterman, E.J.G. (2017). Ensemble and single-molecule dynamics of IFT dynein in Caenorhabditis elegans cilia. Nat. Commun. 8, 14591.

Moore, J.K., Stuchell-Brereton, M.D., and Cooper, J.A. (2009). Function of dynein in budding yeast: mitotic spindle positioning in a polarized cell. Cell Motil. Cytoskeleton 66, 546–555.

Nicholas, M.P., Berger, F., Rao, L., Brenner, S., Cho, C., and Gennerich, A. (2015). Cytoplasmic dynein regulates its attachment to microtubules via nucleotide state-switched mechanosensing at multiple AAA domains. Proc. Natl. Acad. Sci. U. S. A. 112, 6371–6376.

Niekamp, S., Coudray, N., Zhang, N., Vale, R.D., and Bhabha, G. (2019). Coupling of ATPase activity, microtubule binding, and mechanics in the dynein motor domain. EMBO J. 38, e101414.

Pisa, R., Cupido, T., and Kapoor, T.M. (2019). Designing Allele-Specific Inhibitors of Spastin, a Microtubule-Severing AAA Protein. J. Am. Chem. Soc. 141, 5602–5606.

Pöhler, R., Krahn, J.H., van den Boom, J., Dobrynin, G., Kaschani, F., Eggenweiler, H.-M., Zenke, F.T., Kaiser, M., and Meyer, H. (2018). A non-competitive inhibitor of VCP/p97 and VPS4 reveals conserved allosteric circuits in type I and II AAA ATPases. Angew. Chem. Int. Ed. 57, 1576–1580.

Pomeroy, S.L., Tamayo, P., Gaasenbeek, M., Sturla, L.M., Angelo, M., McLaughlin, M.E., Kim, J.Y.H., Goumnerova, L.C., Black, P.M., Lau, C., et al. (2002). Prediction of central nervous system embryonal tumour outcome based on gene expression. Nature 415, 436–442.

Reck-Peterson, S.L., Yildiz, A., Carter, A.P., Gennerich, A., Zhang, N., and Vale, R.D. (2006). Single-molecule analysis of dynein processivity and stepping behavior. Cell 126, 335–348.

Roberts, A.J. (2018). Emerging mechanisms of dynein transport in the cytoplasm versus the cilium. Biochem. Soc. Trans. 46, 967–982.

Robertson, M.J., Van Zundert, G.C.P., Borrelli, K., and Skiniotis, G. (2019). GemSpot: A Pipeline for Robust Modeling of Ligands into CryoEM Maps. BioRxiv.

Rohou, A., and Grigorieff, N. (2015). CTFFIND4: Fast and accurate defocus estimation from electron micrographs. J. Struct. Biol. 192, 216–221.

Ruhnow, F., Zwicker, D., and Diez, S. (2011). Tracking single particles and elongated filaments with nanometer precision. Biophys. J. 100, 2820–2828.

Schmidt, H., and Carter, A.P. (2016). Review: Structure and mechanism of the dynein motor ATPase. Biopolymers 105, 557–567.

Schmidt, H., Gleave, E.S., and Carter, A.P. (2012). Insights into dynein motor domain function from a 3.3-Å crystal structure. Nat. Struct. Mol. Biol. 19, 492–497, S1.

Schmidt, H., Zalyte, R., Urnavicius, L., and Carter, A.P. (2015). Structure of human cytoplasmic dynein-2 primed for its power stroke. Nature 518, 435–438.

See, S.K., Hoogendoorn, S., Chung, A.H., Ye, F., Steinman, J.B., Sakata-Kato, T., Miller, R.M., Cupido, T., Zalyte, R., Carter, A.P., et al. (2016). Cytoplasmic Dynein Antagonists with Improved Potency and Isoform Selectivity. ACS Chem. Biol. 11, 53–60.

Steinman, J.B., and Kapoor, T.M. (2019). Using chemical inhibitors to probe AAA protein conformational dynamics and cellular functions. Curr. Opin. Chem. Biol. 50, 45–54.

Steinman, J.B., Santarossa, C.C., Miller, R.M., Yu, L.S., Serpinskaya, A.S., Furukawa, H., Morimoto, S., Tanaka, Y., Nishitani, M., Asano, M., et al. (2017). Chemical structure-guided design of dynapyrazoles, cell-permeable dynein inhibitors with a unique mode of action. Elife 6.

Subramanian, R., Ti, S.-C., Tan, L., Darst, S.A., and Kapoor, T.M. (2013). Marking and measuring single microtubules by PRC1 and kinesin-4. Cell 154, 377–390.

Tang, W.K., Odzorig, T., Jin, W., and Xia, D. (2019). Structural Basis of p97 Inhibition by the Site-Selective Anticancer Compound CB-5083. Mol. Pharmacol. 95, 286–293.

Thompson, A.R., Hoeprich, G.J., and Berger, C.L. (2013). Single-molecule motility: statistical analysis and the effects of track length on quantification of processive motion. Biophys. J. 104, 2651–2661.

Thorn, K.S., Ubersax, J.A., and Vale, R.D. (2000). Engineering the processive run length of the kinesin motor. J. Cell Biol. 151, 1093–1100.

Ti, S.-C., Pamula, M.C., Howes, S.C., Duellberg, C., Cade, N.I., Kleiner, R.E., Forth, S., Surrey, T., Nogales, E., and Kapoor, T.M. (2016). Mutations in Human Tubulin Proximal to the Kinesin-Binding Site Alter Dynamic Instability at Microtubule Plus- and Minus-Ends. Dev. Cell 37, 72–84.

Toropova, K., Zou, S., Roberts, A.J., Redwine, W.B., Goodman, B.S., Reck-Peterson, S.L., and Leschziner, A.E. (2014). Lis1 regulates dynein by sterically blocking its mechanochemical cycle. eLife 3.

Zhang, K., Foster, H.E., Rondelet, A., Lacey, S.E., Bahi-Buisson, N., Bird, A.W., and Carter, A.P. (2017). Cryo-EM Reveals How Human Cytoplasmic Dynein Is Autoinhibited and Activated. Cell 769, 1303–1314.e18.

