## Supplementary figures for "Targeting Allostery in the Dynein Motor Domain with Small Molecule Inhibitors"

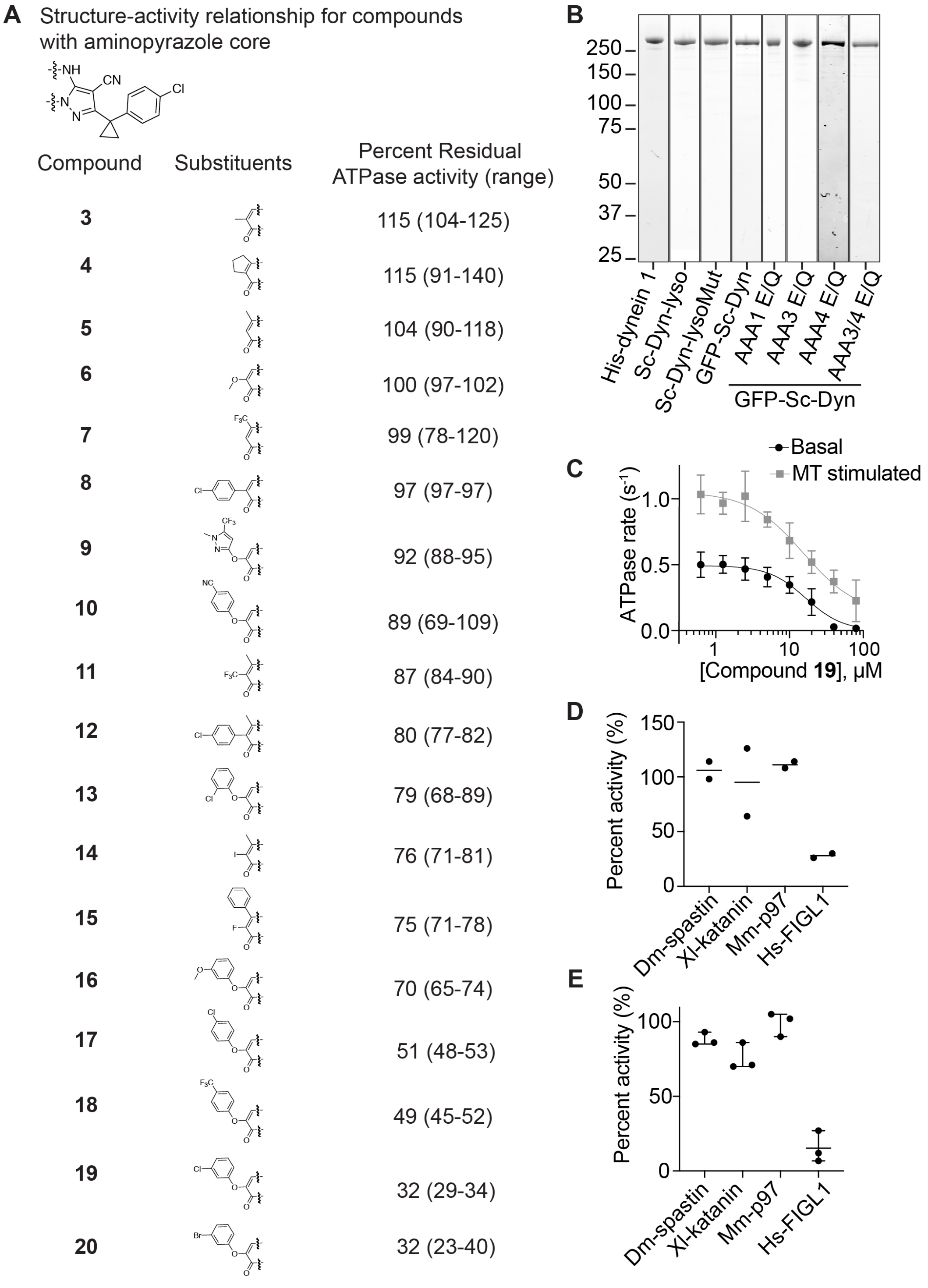


**Figure S1. Structure-activity relationship studies of dynapyrazole derivatives.** (A) Basal ATPase activity of Hs-dynein 1 in the presence of dynapyrazole derivatives (20 µM). Chemical structure of each dynapyrazole derivative is shown. (B) SDS-PAGE analysis of purified dynein constructs (Coomassie blue). (C) ATPase activity of Hs-dynein 1 in the presence of compound **19** with or without the addition of microtubules (2.5 µM). Data are mean ± SD of n=3 and were fit to a sigmoidal dose-response curve. (D) Percent steady-state ATPase activity of four AAA proteins in the presence of compound **19** (20 µM). Lines represent mean (n=2). Mean for each construct is as follows: 106% (Dm-spastin), 95% (Xl-katanin), 111% (Mm-97), 28% (Hs-FIGL). (E) Percent steady-state ATPase activity of four AAA proteins in the presence of compound **20** (20 µM). Lines represent mean and error bars indicate SD (n=3). Mean ± SD for each construct is as follows: 88 ± 4.4 % (Dm-spastin), 99 ± 7.9 % (Mm-p97), 76 ± 9.0% (Xl-katanin), 15 ± 11% (Hs-FIGL).

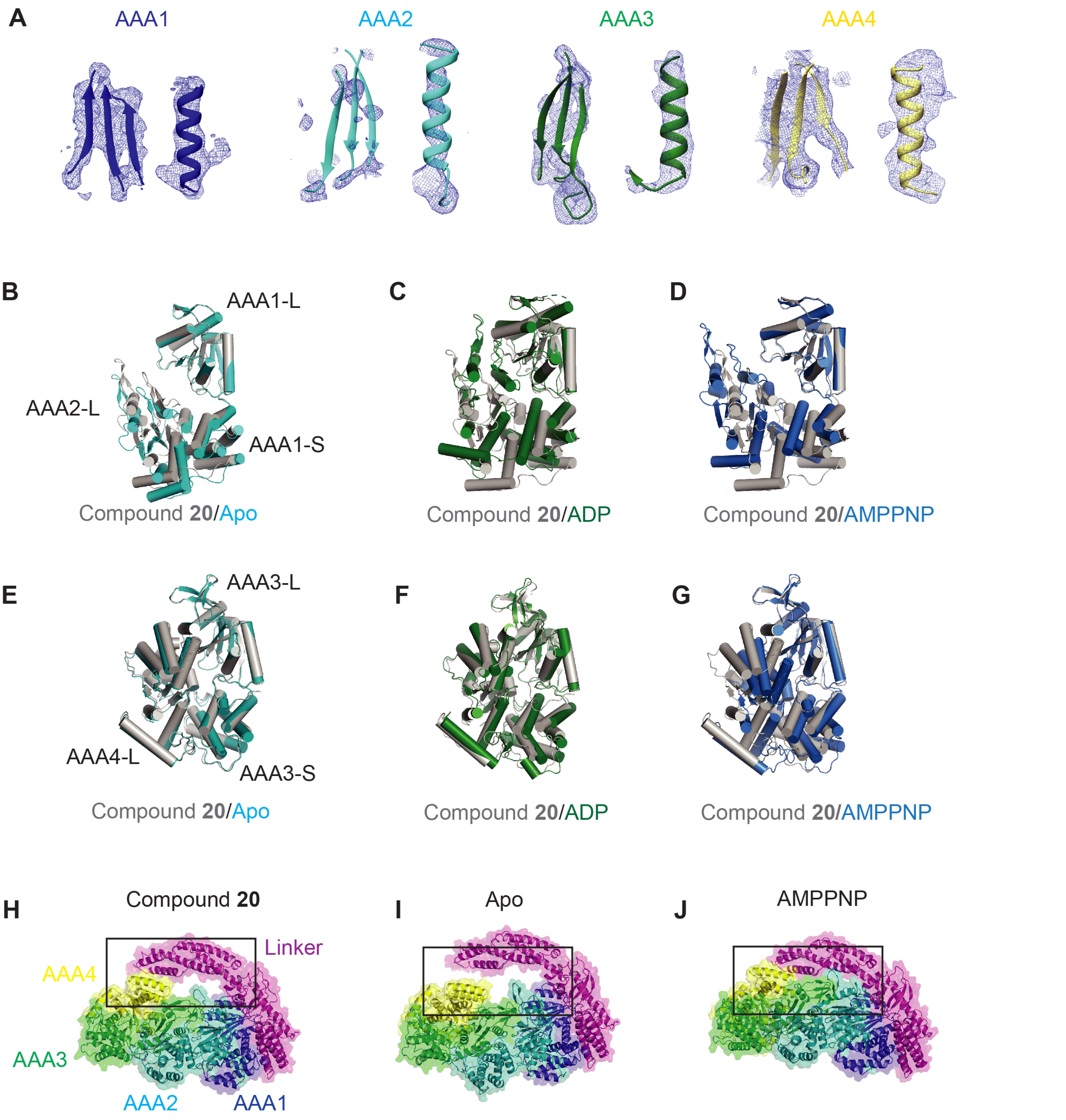


**Figure S2. X-ray model of Sc-Dyn-lysoMut in the presence of compound 20.** (A) Electron densities (blue mesh) of $\beta$-strands and $\alpha$-helices in the AAA1, AAA2, AAA3, and AAA4 sites. Color coding of domains is the same as in Figure 3A. (B-D) Comparison of the AAA1 domain between the X-ray model (gray) and either the *S. cerevisiae* apo (cyan) (B), human ADP (PDB: 5NUG, green) (C), or *S. cerevisiae* AMPPNP (PDB: 4W8F, blue) (D) model. Models are aligned on the AAA1-L subdomain. (E-G) Comparison of the AAA3 domain between the X-ray model (gray) and either the *S. cerevisiae* apo (cyan) (E), human ADP (PDP: 5NUG, green) (F), or Sc-Dyn-lysoMut AMPPNP (PDB: 4W8F, blue) (G) model. Models are aligned on the AAA3-L subdomain. (H-J) Comparison of one side of the AAA ring in the X-ray model (H), apo-model (I), and the AMPPNP-model (J) (PDB: 4W8F). The box highlights the gap between the linker and the AAA1/AAA2/AAA3/AAA4 domains. Color coding of domains is the same as in Figure 3A; models are aligned on the AAA1-L subdomain.

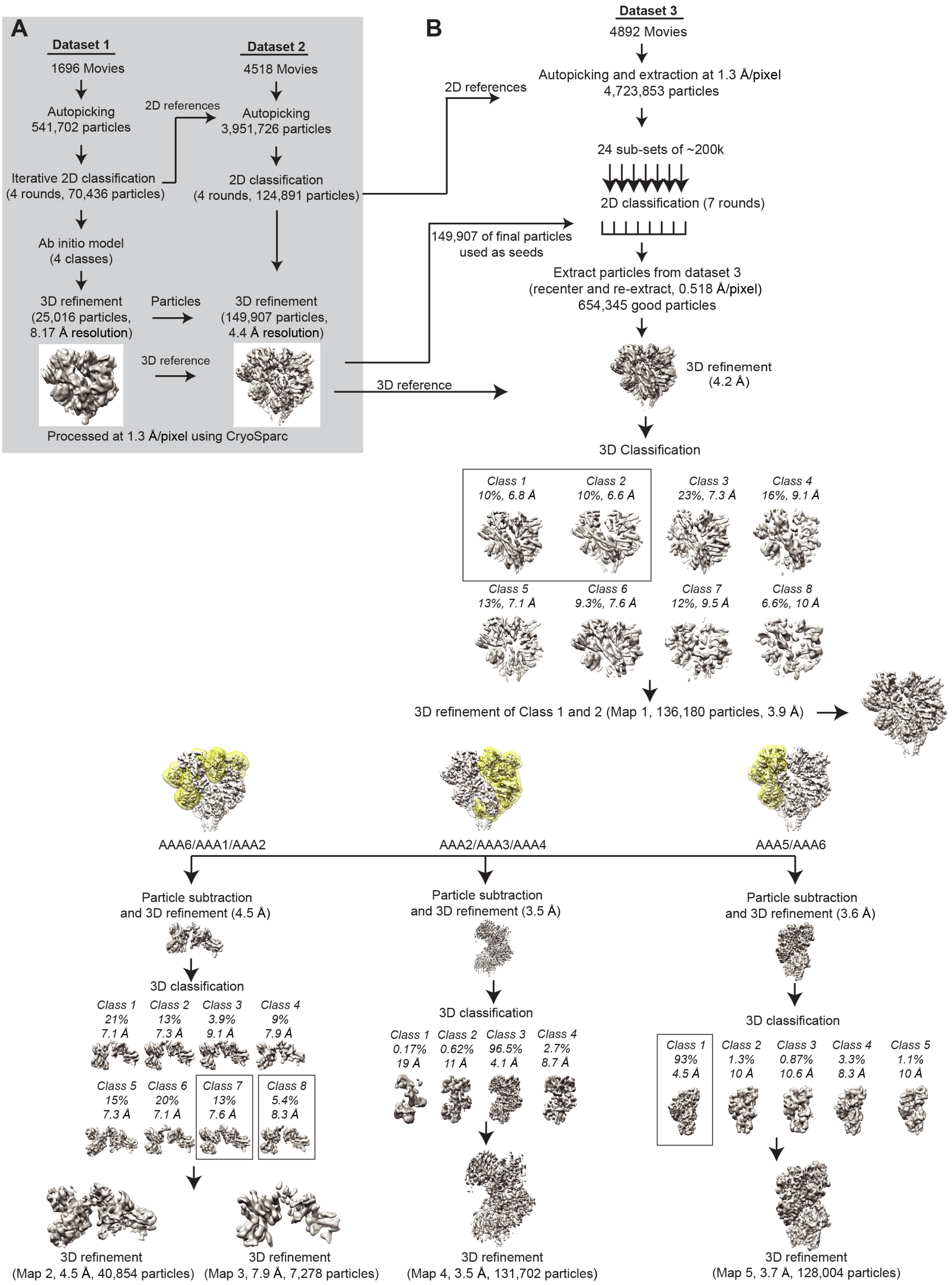


**Figure S3. Processing of the Cryo-EM data for Sc-Dyn-lysoMut in the presence of compound 20.** (A) The workflow used to process datasets 1 and 2 in CryoSparc. (B) The workflow used to process dataset 3 in RELION 3.0.

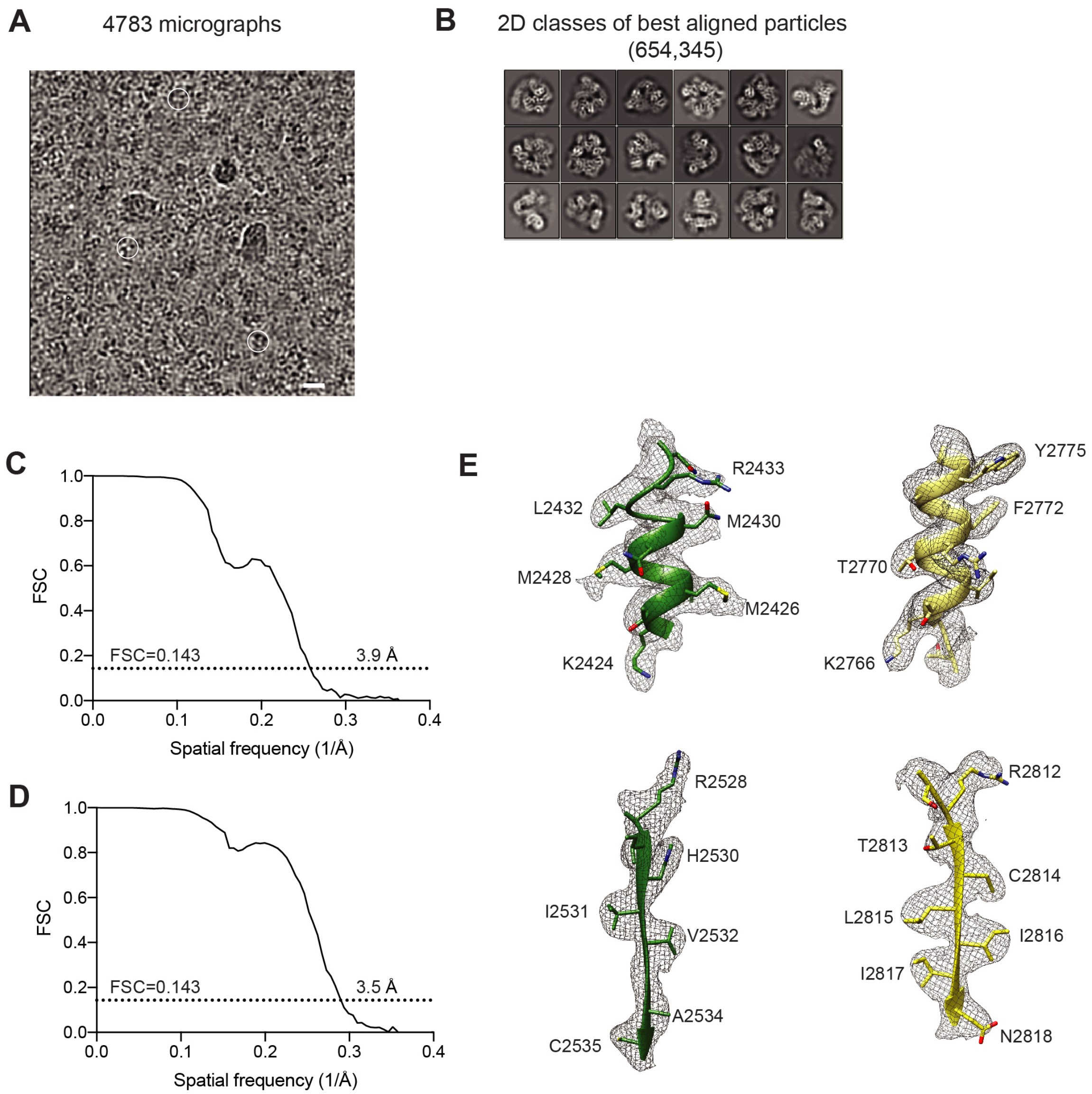


**Figure S4. Cryo-EM analysis of Sc-Dyn-lysoMut in the presence of compound 20.** (A) Representative micrograph from dataset 3 (total: 4892 micrographs). White scale bar represents 20 nm. (B) 2D classes of best aligned particles (654,345). (C) Gold-standard Fourier Shell Correlation (FSC) curve calculated for the Sc-Dyn-lysoMut map (see Figure S3B). The resolution was estimated at ~3.9 Å (FSC=0.143). (D) Gold-standard Fourier Shell Correlation (FSC) curve calculated for the signal subtracted map of the AAA2/AAA3/AAA4 domains (see Figure S3B). The resolution was estimated at ~3.5 Å (FSC=0.143). (E) EM densities (Map 4, gray mesh) of $\beta$-strands and $\alpha$-helices in the AAA3 (green) and AAA4 (yellow) domains. Amino acid residues are indicated and the color coding of domains is the same as in Figure 3A.

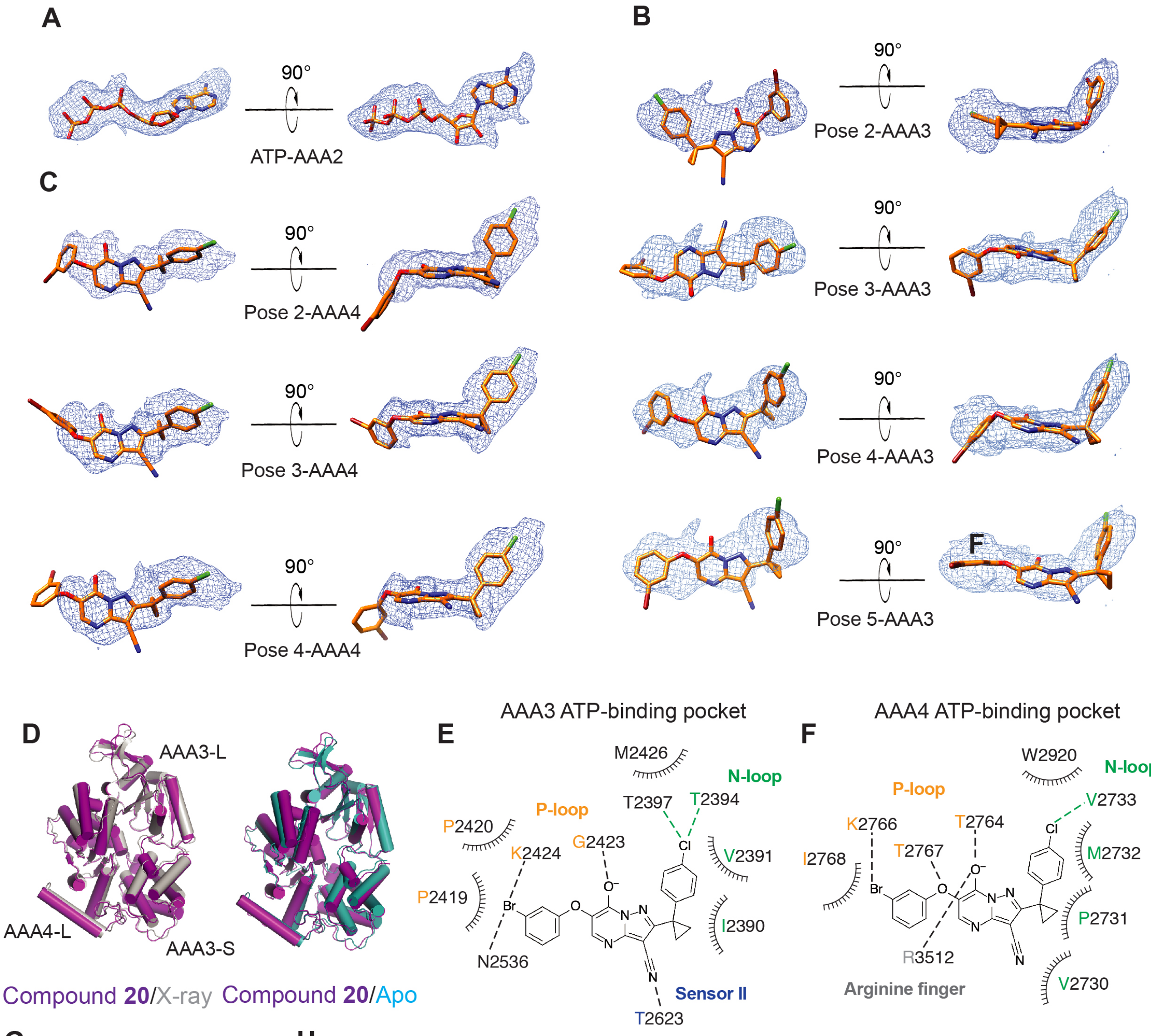


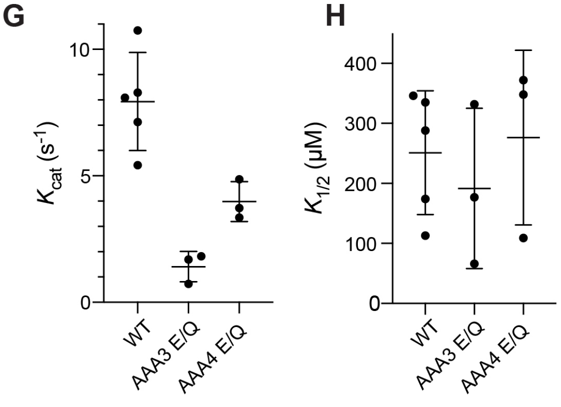


I

| GFP-Sc-Dyn | $k_{cat}$ ($s^{-1}$) | $k_{1/2}$ (µM) | Hill coefficient |
|---|---|---|---|
| WT | 7.9 (1.9) | 251 (103) | 0.81 (0.10) |
| AAA3 E/Q | 1.4 (0.6) | 192 (133) | 0.36 (0.05) |
| AAA4 E/Q | 4.0 (0.8) | 276 (145) | 0.77 (0.33) |

**Figure S5. Analysis of compound binding to Sc-Dyn-lysoMut.** (A) Orientation of ATP in the AAA2 site generated by the GlideEM script and overlayed with the EM density (Map 4, blue mesh). ATP is shown as a stick model (carbon: orange, oxygen: red, nitrogen: blue). (B, C) Poses of compound **20** generated by the Glide EM script for the AAA3 (B) and AAA4 (C) sites and overlayed with the EM density (Map 4, blue mesh). Compound **20** is shown as a stick model (carbon: orange, oxygen: red, nitrogen: blue, chlorine: green, bromine: dark red). (D) Comparison of the AAA3 domain between the cryo-EM reconstruction of Map 3 (magenta) and either the X-ray model (gray) or apo-model (cyan). Models are aligned on the AAA3-L subdomain. (E, F) Schematic for the predicted hydrogen bonding (black dashed lines), halogen bonding (green dashed lines) and van der waals interactions (ticked curved lines) between compound **20** and either the N-loop, P-loop or sensor II motifs (N-loop: green, P-loop: yellow, sensor II: blue) in the AAA3 (E) or AAA4 (F) nucleotide-binding pockets. (G, H) Catalytic turnover number ($k_{cat}$; G) and ATP concentration required for half-maximal velocity ($k_{1/2}$; H) of GFP-Sc-Dyn and its mutants. Data represent average ± s.d. (n≥3). Basal ATPase activity of the double mutant (AAA3/AAA4 E/Q) was too weak to accurately determine the construct's enzymatic parameters. (I) Values for enzymatic activity parameters of GFP-Sc-Dyn and its mutants are provided (mean, SD in parentheses).

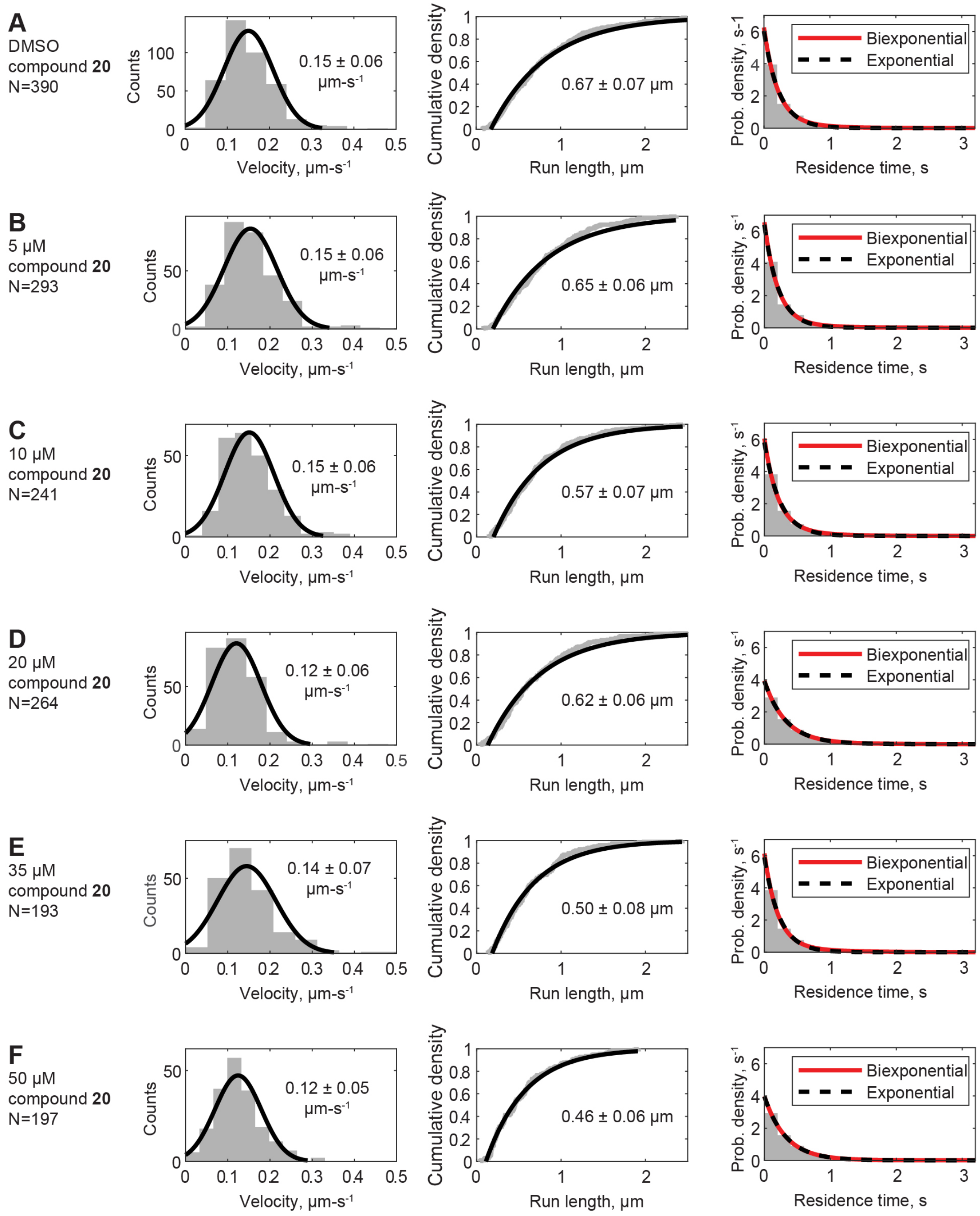


**Figure S6. Effect of compound 20 on the velocity, run length, and pause density of Sc-DynGST.** (A-F) All data are pooled from n=3-4 independent experiments, with the number of events depicted in each panel. Velocities are shown with fit to a normal distribution, data reported as mean ± SD. Run length distributions are shown as cumulative density function with offset exponential fit. Rightmost column of data shows the probability distribution of residence times of individual Sc-DynGST molecules within 50-nm bins along the direction of motion, following a published method (DeWitt et al., 2015). Fitting to single exponentials (black dashed lines) and bi-exponentials (red lines) yielded nearly identical results.

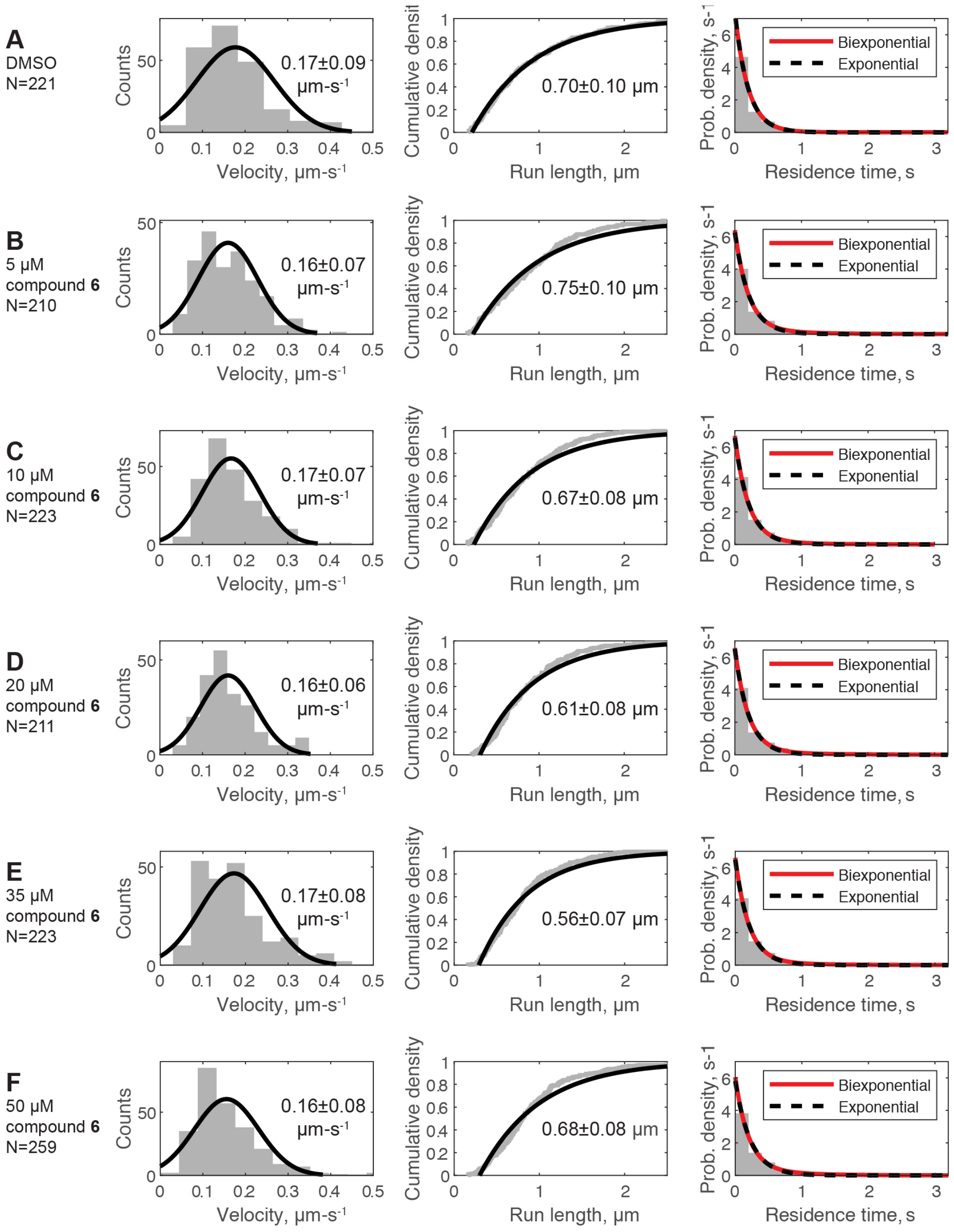


**Figure S7. Effect of compound 6 on the velocity, run length, and pause density of Sc-DynGST.** (A-F) All data are pooled from n=3 independent experiments, with the number of events depicted in each panel. Velocities are shown with fit to a normal distribution, data reported as mean ± SD. Run length distributions are shown as cumulative density function with offset exponential fit. Rightmost column of data shows the probability distribution of residence times of individual Sc-DynGST molecules within 50-nm bins along the direction of motion, following a published method (DeWitt et al., 2015). Fitting to single exponentials (black dashed lines) and bi-exponentials (red lines) yielded nearly identical results.
