## Supplementary material for "Targeting Allostery in the Dynein Motor Domain with Small Molecule Inhibitors": Table S1

**Table S1** Data collection and refinement statistics (molecular replacement)

|  | Sc-Dyn-lysoMut |
| --- | --- |
| **Data collection** |  |
| Space group | *P2_1_2_1_2* |
| Cell dimensions |  |
| *a*, *b*, *c* (Å) | 135.06, 157.92, 179.31 |
| α, β, γ (°) | 90.00 90.00 90.00 |
| Resolution (Å) | 47.66-4.50 (4.77-4.50) |
| *R*_meas_ | 28.6 (168.7) |
| *I* / σ*I* | 5.99 (1.13) |
| Completeness (%) | 99.5 (98.3) |
| Redundancy | 6.8 (6.5) |
| CC1/2 | 48.4 |
| **Refinement** |  |
| Resolution (Å) | 50-4.5 |
| No. reflections | 157857 |
| *R*_work_ / *R*_free_ | 0.247/0.289 |
| No. atoms |  |
| Protein | 42311 |
| Ligand/ion | N/A |
| Water | N/A |
| *B*-factors |  |
| Protein | 225 |
| Ligand/ion | N/A |
| Water | N/A |
| R.m.s. deviations |  |
| Bond lengths (Å) | 0.002 |
| Bond angles (°) | 0.5 |
| Clashscore | 5.3 |

Values are for a single crystal.

*Highest-resolution shell is shown in parentheses.
