## Supplementary material for "Targeting Allostery in the Dynein Motor Domain with Small Molecule Inhibitors": Table S2

| **Data Collection** | Dataset 1 | | Dataset 2 | | Dataset 3 |
| --- | --- | --- | --- | --- | --- |
| Microscope | Titan Krios | | Titan Krios | | Titan Krios |
| Camera | K2 Summit | | K2 Summit | | K2 Summit |
| Voltage (kV) | 300 | | 300 | | 300 |
| Frames | 50 | | 50 | | 50 |
| Exposure time (s) | 10 | | 10 | | 10 |
| Total dose (ē/Å^2^) | 71 | | 52 | | 44 |
| Defocus range | -1.5 to -3 | | -0.9 to -1.7 | | -1 to -1.5 |
| Super resolution pixel size (Å) | 0.6675 | | 0.6675 | | 0.518 |
| Micrographs | 1696 | | 4518 | | 4893 |
| **Model composition** | Map 1 | Map 2 | Map 3 | Map 4 | Map 5 |
| Non-hydrogen atoms | 39189 | 14794 | 14794 | 19070 | 14430 |
| Protein residues | 2414 | 936 | 936 | 677 | 889 |
| Ligands (ATP/Compound **19**) | 1/2 | 0/0 | 0/0 | 1/2 | 0/0 |
| Resolution | ~3.9 | ~4.7 | ~7.9 | ~3.5 | ~3.7 |
| Map sharpening B-factors (Å2) | -100 | -100 | -50 | -50 | -100 |
| Overall FSC† | 0.143 | 0.143 | 0.143 | 0.143 | 0.143 |
| Correlation coefficient | 0.74 | 0.63 | 0.52 | 0.85 | 0.77 |
| Mean B-factor (Å^2^) | 89.31 | 106.89 | 287.87 | 139.64 | 80.18 |
| rmsd (bonds) | 0.015 | 0.016 | 0.017 | 0.013 | 0.011 |
| rmsd (angles) | 1.397 | 0.982 | 0.912 | 1.454 | 1.125 |
| MolProbity score | 2.76 | 2.17 | 2.40 | 2.34 | 2.19 |
| Clashscore, all atoms | 26.68 | 6.09 | 10.89 | 15.32 | 10.90 |
| Rotamer outliers | 5.34 | 3.49 | 3.49 | 2.03 | 2.2 |
| Ramachandran plot: |  |  |  |  |  |
| Favored | 94.09 | 93.6 | 93.44 | 92.59 | 95.60 |
| Outliers | 0.46 | 0.54 | 0.65 | 1.68 | 0.46 |

**Table S2** Cryo-EM data collection and refinement statistics
